# A potent myeloid response is rapidly activated in the lungs of premature Rhesus macaques exposed to intra-uterine inflammation

**DOI:** 10.1101/2021.05.28.444219

**Authors:** Courtney M. Jackson, Martin Demmert, Shibabrata Mukherjee, Travis Isaacs, Jerilyn Gray, Paranthaman Senthamaraikannan, Pietro Presicce, Kashish Chetal, Nathan Salomonis, Lisa A. Miller, Alan H. Jobe, Suhas G. Kallapur, William J. Zacharias, Ian P. Lewkowich, Hitesh Deshmukh, Claire A. Chougnet

## Abstract

Intrauterine inflammation/infection (IUI), which is present in up to 40% of premature births, leads to elevated levels of pro-inflammatory mediators and microbial products within the amniotic fluid, which come in close contact to fetal mucosae. Yet, knowledge on the fetal mucosal responses to IUI exposure remains limited. To address these questions, we used a non-human primate model of IUI, in which pregnant Rhesus macaques received intra-amniotic (IA) LPS, compared with IA saline.

We found that IA LPS exposure induced a robust and rapid inflammation of the fetal lung, but not the intestine. This inflammatory response was characterized by high levels of pro-inflammatory cytokines in the lung and the alveolar wash, and a potent myeloid cell response, dominated by neutrophils and monocytes/macrophages. scRNAseq analyses of fetal lungs showed that the infiltrating (neutrophils and inflammatory monocytes) and the resident (alveolar and interstitial macrophages) myeloid cells exhibited transcriptional profiles consistent with exposure to TLR ligands, as well as to cytokines, notably IL-1 and TNFα. However, blocking IL-1 signaling or TNFα, alone or simultaneously by administering inhibitors intra-amniotically and subcutaneously to the dam only partially blunted fetal lung inflammation.

Together, our novel data indicate that the fetal innate immune system can mount a rapid multi-factorial mucosal innate response to IUI, responding both to direct signaling by bacterial products and to indirect cytokine-mediated pathways of activation. These data thus provide more mechanistic insights into the association between IUI exposure and the post-natal lung morbidities of the premature infant.

## Introduction

Intrauterine inflammation/infection (IUI) is present in 25-40% of premature births^1^. Inflammation of the placental membranes at the fetal-maternal interface, which is called chorioamnionitis (chorio)^2^, leads to elevated levels of pro-inflammatory mediators and microbial products within the amniotic fluid ^2–5^ Due to *in utero* aspiration/swallowing of the amniotic fluid ^6,7^, the mucosae of the developing fetus come in close contact with these mediators, which triggers an inflammatory response. Although such responses likely contribute to the increased incidence of post-natal pulmonary and intestinal morbidities associated with IUI exposure ^8–10^, knowledge about the cells and pathways involved in the fetal mucosal response remains very limited, due to the inability to collect tissue samples in preterm infants.

Fetal lung and intestinal inflammation have been described in IUI animal models (rodent, rabbits, pig, and sheep). Main features include high levels of pro-inflammatory cytokines and infiltration of immune cells, as well as damage to the lung and intestine ^11–18^. However, the ontogeny of the immune system or the placenta structure are quite different in these species from those in humans. In contrast, non-human primates (NHP) are quite similar to humans in most biological aspects. Furthermore, many reagents developed against human antigens are cross-reactive allowing for in-depth analysis of the NHP fetal immune system^19–21^. Importantly, we have previously shown that intra-amniotic (IA) injection of LPS in pregnant Rhesus macaques leads to a type of inflammation that closely phenocopies severe human chorio ^22,23^. Previous studies from ours and other groups have reported elevated levels of inflammatory cytokines in the lung of IUI-exposed rhesus macaque fetuses ^24,23^, but no in-depth analysis of mucosal immune responses was performed in these studies. Our goal herein was thus to apply state-of-the art technologies, such as single-cell RNAseq (scRNAseq), to obtain a comprehensive view of the mucosal immune responses occurring in response to IUI.

Furthermore, this model allows us to evaluate the role inflammatory cytokines may play in these responses. Among the cytokines elevated in the amniotic fluid and cord blood in chorio, IL-1β and TNFα appear particularly important ^4,25^. Indeed, both cytokines are capable of inducing chorio, and neutrophil infiltration into the fetal lung of Rhesus macaques when administered IA to pregnant dams ^26,27^. In addition, IL-1β is involved in many aspects of LPS-induced fetal inflammation ^22,23,27–30^. We thus aimed at better delineating their respective role, using the cross-reactive IL-1 receptor antagonist (IL-1RA) and a blocking anti-TNF Ab (Adalimumab) given either individually, or in combination.

Herein, we show that IA LPS leads to rapid and robust inflammation of the fetal lung, but not of the fetal intestine. Notably, our analyses identified two main components, i.e., the recruitment of innate myeloid cells into the fetal lung, and the activation of resident lung myeloid cells. Although signaling by IL-1 and TNFα were identified as major transcriptional signatures in responsive myeloid cells, their individual or combined blockade only partially blunted fetal lung inflammation. Together, our novel data thus indicate that the fetal innate immune system is capable of mounting a rapid multi-factorial mucosal response to IUI, responding both to direct signaling by bacterial products such as LPS, and to indirect cytokine-mediated pathways of activation.

## Results

### IA LPS induces robust inflammation in the fetal lung with a minimal response in the small intestine

To model IUI, pregnant Rhesus macaques were given intra-amniotic (IA) injection of LPS or saline at approximately 80% gestation. Sixteen hours post IA injections, the fetuses were surgically removed, and tissues collected for analysis. All the animals included in the study are described in Table S1.

We first examined the histological changes in the fetal lung and small intestine. Compared to controls, lungs from 16hr IA LPS-exposed fetuses displayed signs of inflammation with cell infiltration into the lung interstitium and decreased alveolar space (Fig. 1A). Secondary alveoli septa formation also appeared diminished (Fig. 1A). In contrast, in the jejunum of exposed fetuses, the intestinal villi, lamina propria, or the underlying mucosal area exhibited no abnormalities (Fig. S1A). Because intestinal inflammation did not develop until 3 days after IUI in other models (pig^18^ and sheep ^12,17,31^), we analyzed jejunum sections taken 48 and 120hrs post IA LPS, and again, found only minimal changes compared to controls (Fig. S1A).

**Fig. 1.**
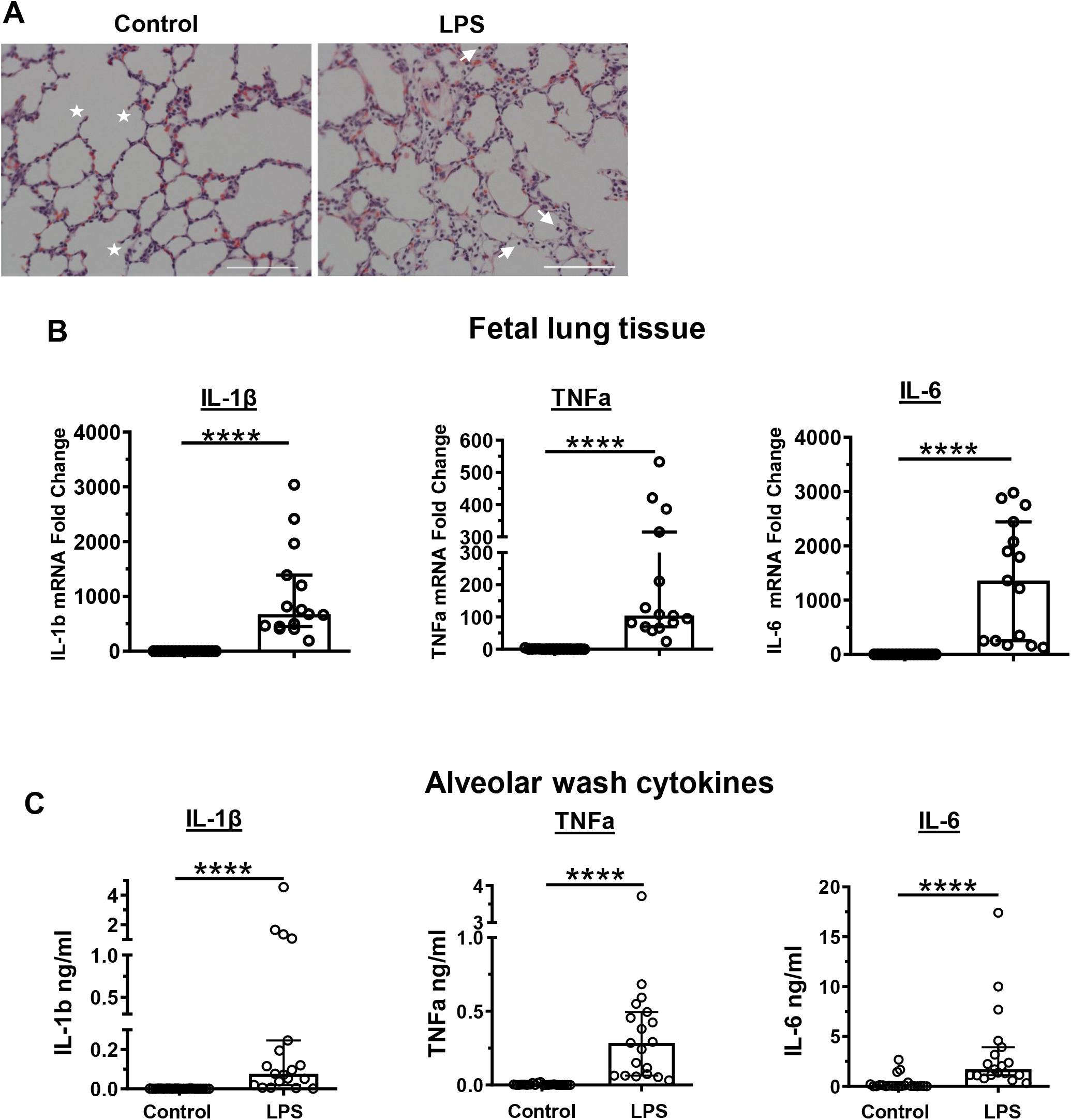
Fetal lung response 16 hours post IA LPS exposure. **(A)** 20x hematoxylin & eosin (H&E) stain of fetal lung from control and LPS exposed fetuses at 16 hours following IA LPS exposure; stars denote secondary septa, arrows represent infiltrating cells in alveolar space and scale bar is 100μm. IL-1β, TNFα, and IL-6, mRNA expression in fetal lung tissue **(B)** and concentration in the alveolar wash **(C)** in control and 16 hours post IA LPS. Data presented as median and interquartile range, Mann-Whitney U test; ****p≤0.0001.

Along with histological evaluations, mRNA expression of inflammatory cytokines was measured in lung and jejunum at 16hrs post IA LPS. TNFα, IL-6, and IL-1β mRNA levels were greatly increased in the lung (>100-fold) of IA LPS-exposed fetuses along with increased detection of these cytokines in the alveolar wash (AW) (Fig. 1B, C). Other pro-inflammatory cytokines such as IL-8 and CCL2 were also increased (see Tables S2 and S3). In contrast, IL-10, a known anti-inflammatory molecule was only minimally increased in the AW of LPS-exposed infants (Table S2). In agreement with the absence of histological alterations in the jejunum, mRNA expression of inflammatory cytokines was not significantly increased in this compartment (Fig. S1B). Thus, IA LPS induces a rapid inflammatory response within the fetal lung but has minimal impact on the fetal small intestine. Our in-depth analyses of fetal mucosal responses thus focused on the lung.

### IA LPS triggers a large infiltration of myeloid cells into the fetal lung

We next evaluated by scRNAseq the global transcriptional response of the lung in response to IA LPS (animals and sequencing metrics are described in Table S4). These analyses revealed distinct immune and non-immune cell populations in both control and IA LPS-exposed animals (Fig. 2A, Table S5). As our focus was on immune responses, subsequent analyses were performed on a subset of the whole dataset, containing only the immune cell clusters. (Fig. 2B). Using established lineage marker genes (see Fig. 2C), we identified the robust accrual of neutrophilic and monocyte/macrophage populations in the lung of IA LPS-exposed fetuses, compared to control animals (Fig. 2B). As expected, they highly expressed the myeloid-associated CD88 gene (*C5AR1*, Fig. 2D). These findings were confirmed by flow cytometric analyses of lung cell suspensions (see gating strategy in Fig. S2), which showed a trend toward a higher proportion of CD45^+^ hematopoietic cells, and, within this population, a significant increase in the proportion of CD88^+^ cells (Fig. 2E).

**Fig. 2.**
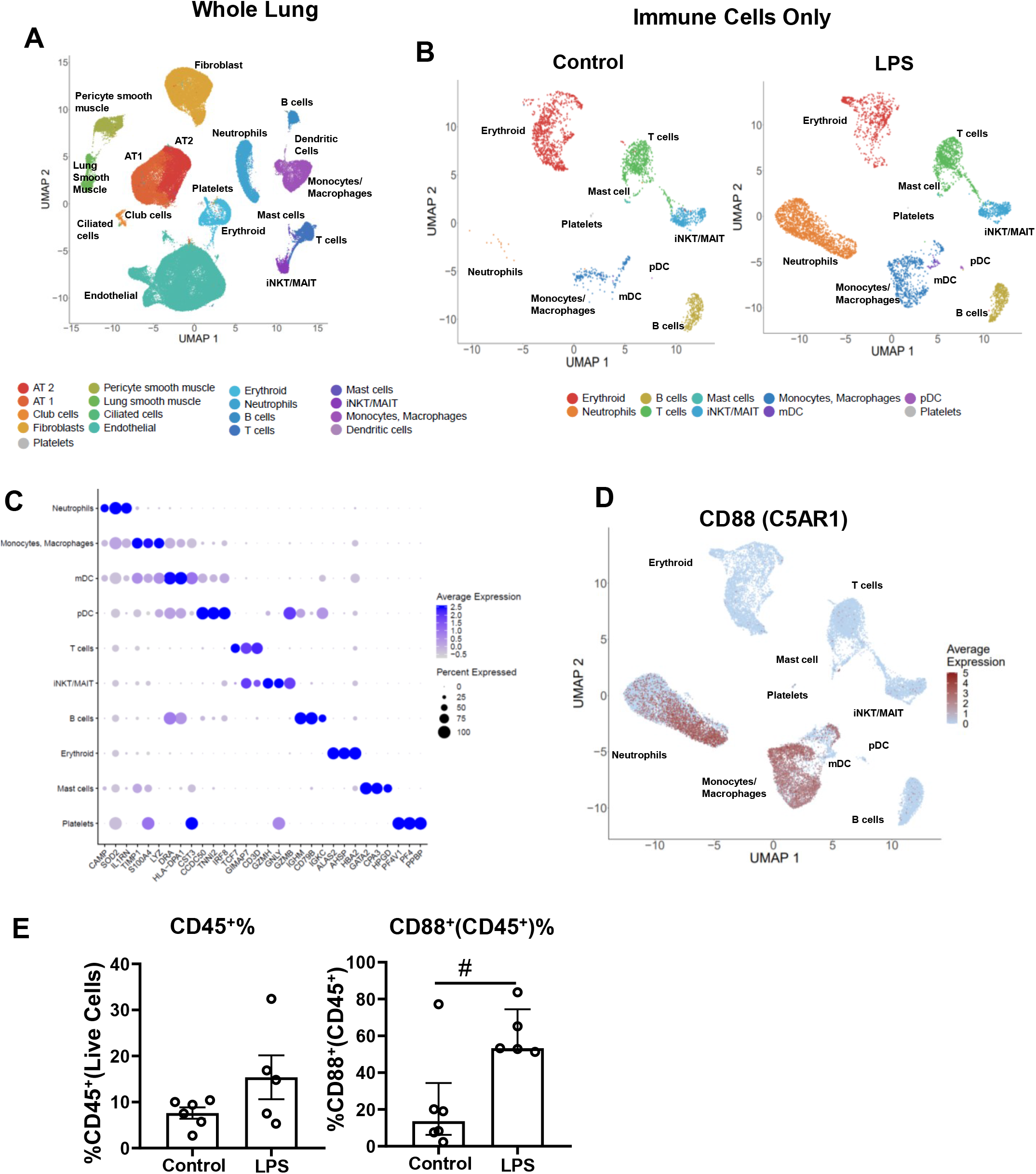
Single cell RNAseq analysis of the fetal lung 16 hours post IA LPS exposure. The left lobe of fetal lung was processed into single cell suspensions and submitted for single cell sequencing. **(A)** UMAP from combined control and LPS-exposed animals (2 each). **(B)** Control (left) and IA LPS (right) exposed animals UMAP of re-clustered immune cell populations. **(C)** Representative dot plot of canonical cell type markers used in the identification of hematopoietic populations. **(D)** Feature plot of CD88(C5AR1) expression across hematopoietic cell populations and **(E)** percentage of total CD45^+^ within live cells and CD88^+^ within CD45^+^ cells in lung. Data presented as mean with SEM, student’s unpaired t-test (CD45^+^%) or median and interquartile range, Mann-Whitney U test (CD88^+^%); #0.05≥p≤0.10.

Due to the rapid response within the lung and the type of challenge used, another myeloid population of interest were dendritic cells (DCs), as DC populations have been described in the human fetal lung^32^. By flow cytometry, we readily identified myeloid dendritic cells (mDCs) and plasmacytoid DCs (pDCs) in the lung, but their frequencies were unchanged by IA LPS (Fig. S3). However, these DCs appeared activated in IA LPS-exposed fetuses, as they both exhibited significantly increased HLA-DR levels (Fig. S3A-B).

In contrast to the brisk myeloid response, the frequency of B cells or different T cell populations, including regulatory T cells, was not changed by IA LPS as seen by flow cytometry and scRNAseq analyses (Fig. S4A-D). Additionally, analyses of differentially expressed genes (DEGs) also uncovered relatively few changes (n=342) in these populations between IA LPS and control animals (Fig S4B, S4D, Table S6).

Taken together, our data show that IA LPS rapidly affected fetal innate immune cell populations in the lung, mainly altering myeloid cells (neutrophils, monocytes/macrophages, DCs), while having little effect on the major lymphoid cell subsets.

### IA LPS leads to the recruitment of inflammatory monocytes into the fetal lung and activates the resident macrophages

scRNAseq analyses of the lungs of control fetuses revealed the presence of 2 macrophage populations (Fig. 3A), which we identified as alveolar and interstitial macrophages (Fig. 3B and S5). In the lung of IA LPS-exposed fetuses, another population of monocytes/macrophages with characteristics of inflammatory monocytes was also present (Fig. 3A-B and Fig. S5).

**Fig. 3.**
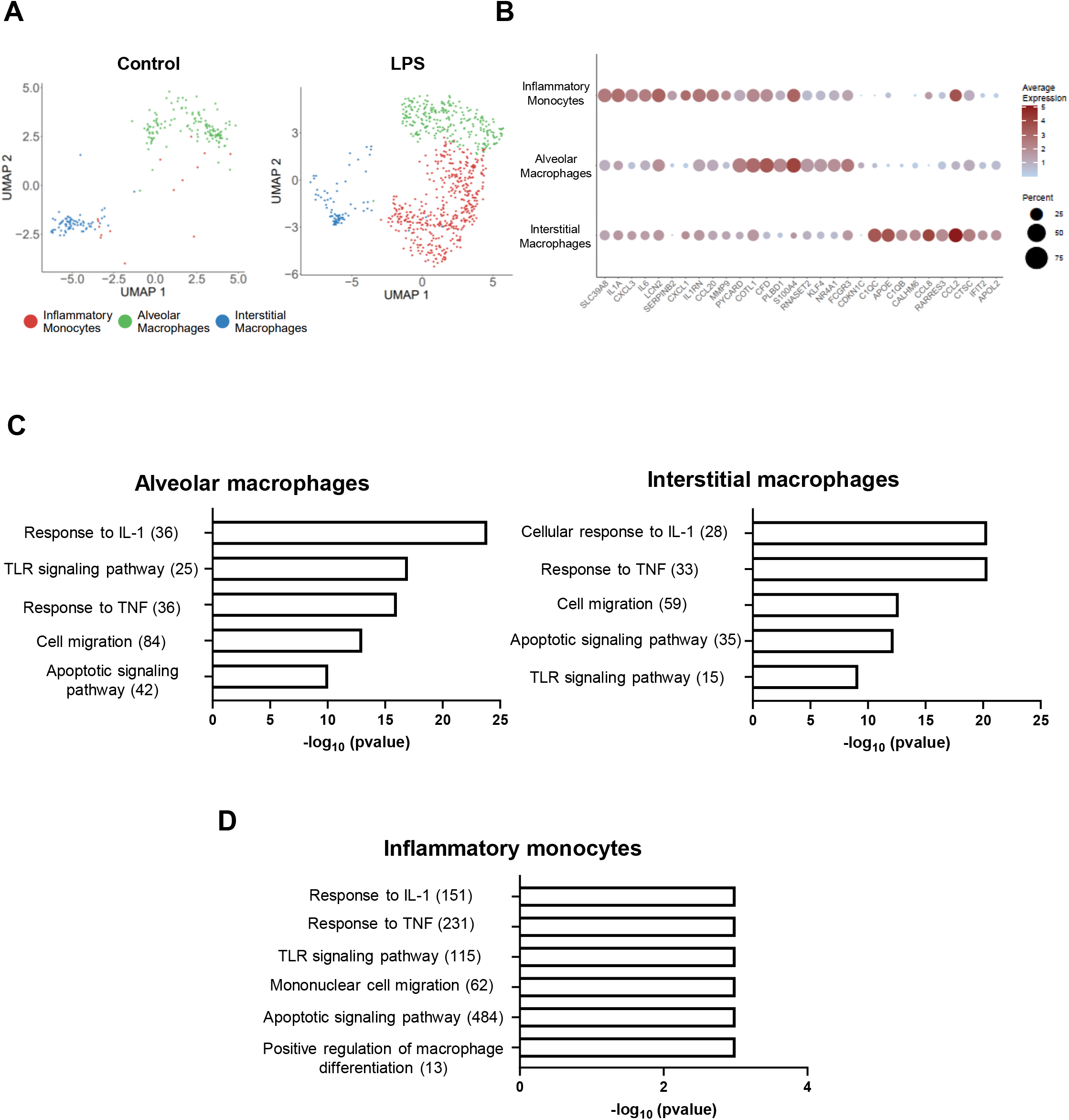
Fetal lung monocyte/macrophage response to IA LPS. **(A)** UMAPs of fetal lung myeloid cell populations in control (left) and IA LPS (right) fetuses. **(B)** Bubble plot of the top 10 conserved genes in the inflammatory monocyte, alveolar macrophage, and interstitial macrophage populations in the fetal lung of IA LPS exposed fetuses. **(C)** Functional enrichment analysis of upregulated biological processes based on differentially expressed genes (alveolar, interstitial macrophages) using ToppFun. **(D)** ssGSEA analysis of enriched gene ontology (GO) terms in inflammatory monocytes. Numbers in parentheses represent the total number of genes for each process.

Next, we identified DEGs in alveolar and interstitial macrophages of IA LPS exposed lungs compared to controls. The majority of these DEGs were more highly expressed in LPS than in controls (n=482/620 in alveolar macrophages and n=302/324 in interstitial macrophages; Table S7, S8). Gene ontology (GO) enrichment analysis of these upregulated genes revealed several biological processes, including activation of the TLR signaling pathway, and response to cytokines, notably to IL-1 and TNF (Fig. 3C, Tables S9 and S10 for full list of genes). Furthermore, genes associated with apoptotic signaling and cell migration were particularly enriched (Fig. 3C). The fewer down-regulated genes in the alveolar and interstitial macrophages of IA LPS lungs mainly included ribosomal protein genes (Tables S6 and S7).

There was no inflammatory monocyte cluster in the control lungs, precluding DEG analysis. However, single sample gene set enrichment analysis (ssGSEA) of this population in the IA LPS-exposed lungs revealed a similar profile as that found in the alveolar and interstitial macrophages from the same lungs. Indeed, there was an enrichment of genes related to activation of the TLR, IL-1 and TNF signaling, as well as cell migration and apoptosis (Fig. 3D, Table S11 for full list of genes expressed and enrichment scores). All together, these data thus show that exposure to IUI strongly affects fetal lung resident macrophages as well as rapidly inducing the afflux of activated, inflammatory monocytes into the fetal lung.

### Recruited neutrophils into the fetal lung have a profile of activation suggesting both direct LPS and indirect cytokine-mediated pathways of activation

Our data also showed a strong neutrophilic response in the fetal lung after IA LPS. scRNAseq analyses identified *TNFAIP6* (TNF Alpha Induced Protein 6) expression as one of the most specific markers for neutrophils (Fig. 4A and Fig. S6A). *S100A8* was also found to be highly expressed in most neutrophils, although it was also present in the monocyte/macrophage cluster (Fig. 4A). The cluster was further uniquely characterized by the expression of neutrophil specific markers *AZU1, ITGAM, NCF2*. By IHC, we confirmed the presence of multiple TNFAIP6^+^S100A8^+^ cells in the IA LPS fetal lungs, which were not present in control lungs (Fig. 4B). These cells had neutrophilic morphology, and they formed aggregates within alveolar spaces (Fig. 4B-C). We confirmed these data by staining with CD68 and HLA-DR, as we had found that CD68 is highly expressed in both fetal neutrophils and monocytes/macrophages, with only the latter expressing HLA-DR. A significantly increased number of CD68^+^HLA-DR^-^ cells, with a neutrophilic morphology, were found in the IA LPS-exposed lungs, also forming aggregates (Fig. S6B). Furthermore, we found a significant number of neutrophils in the AW of IA LPS exposed fetuses, whereas they were largely absent in the AW of control fetuses [median 0.0 v. 106.3×10^6^ cells/kg in control (n=18) and IA-LPS (n=13), respectively, p<0.0001, Mann-Whitney test].

**Fig. 4.**
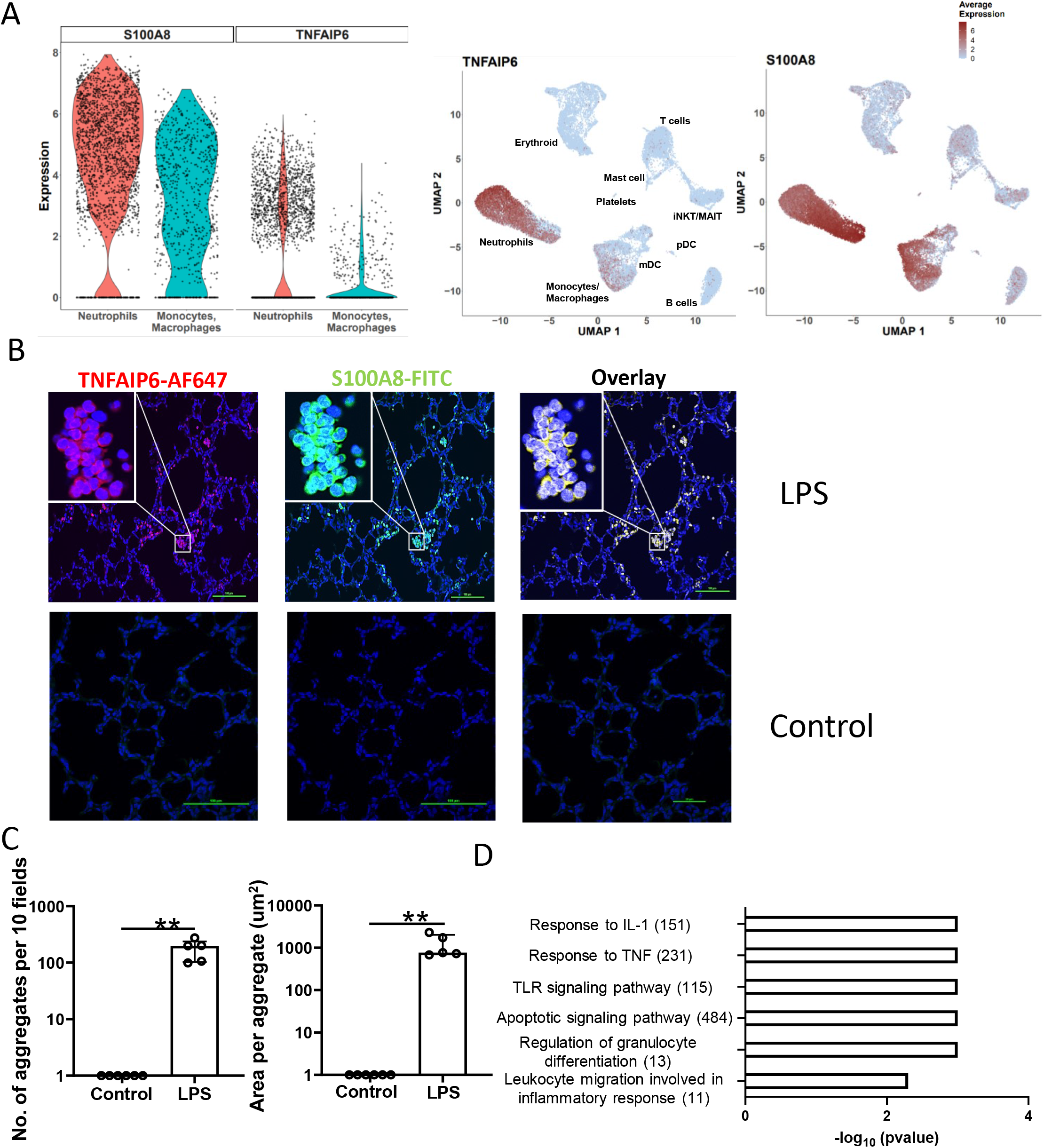
Fetal lung neutrophil response to IA LPS. **(A)** Violin (left) and feature (right) plot of TNFIAP6 and S100A8 expression in fetal lung neutrophils and monocyte/macrophage population following IA LPS exposure. **(B)** Immunofluorescence of IA LPS exposed fetal lung stained for S100A8 and TNFAIP6; scale bar is 100μm. **(C)** Neutrophil aggregate count (left) and area (right) in the fetal lungs of control and IA LPS fetuses. Data presented as median and interquartile range, Mann-Whitney U test; **p≤0.01. **(D)** ssGSEA analysis of enriched GO terms in neutrophils. Numbers in parentheses represent the total number of genes for each process.

We next analyzed the transcriptomic program of the neutrophils recruited in the fetal lungs by ssGSEA. These analyses revealed the enrichment of similar processes as those found in monocytes/macrophages, notably migration, response to TLR, IL-1 and TNF, and apoptosis (Fig.4D and Table S12 for full list of genes expressed and enrichment scores).

Interestingly, and in line with other studies^33,34^, substantial transcriptional heterogeneity could be seen within these recruited neutrophils. However, no distinct sub-clusters were revealed by UMAP projection (Fig. 5A), suggesting different levels of maturation and/or activation. We therefore applied pseudo-time analyses as implemented in Monocle3 to characterize this heterogeneity. Genes that most determined the trajectory were calculated with spatial autocorrelation analysis (Fig. S7). This analysis revealed that cytokine and cytokine-response genes were particularly enriched close to the beginning of the trajectory, whereas expression of other effector genes (e.g., *S100A8/9, CAMP, MMP8*) was higher towards more advanced pseudo-time (Fig. 5, S7 and S8). Thus, the recruited neutrophils into the fetal lung exhibit transcriptional programs revealing different stages of activation, compatible with both direct activation by LPS and indirect cytokine-mediated pathways.

**Fig. 5.**
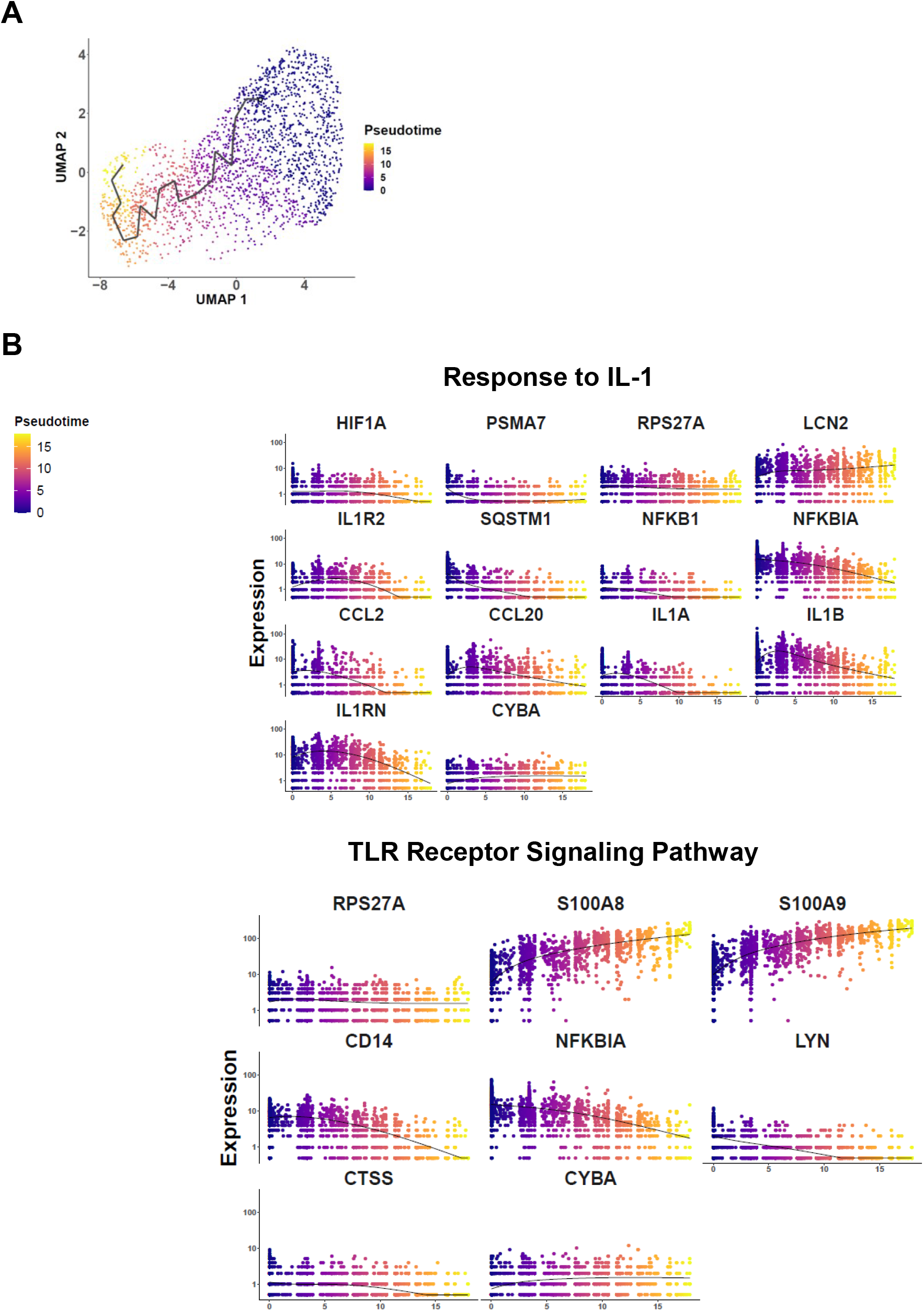
Pseudo-time analysis of fetal lung neutrophil response to IA LPS. **(A)** Pseudo-time trajectory analysis of IA LPS exposed fetal lung neutrophils using Monocle. **(B)** Plots of genes from GO biological pathways response to IL-1 and TLR signaling pathway across pseudo-time.

### IL-1 and TNFα signaling are partially driving fetal lung inflammation

Knowing the role of IL-1 and TNFα in chorio pathogenesis^22,23,26–30^, and because scRNAseq analyses revealed gene signatures consistent with IL-1 and TNFα signaling in both the monocyte/macrophage and neutrophil lung populations of IA LPS animals, we blocked IL-1 signaling with the IL-1RA and TNFα with the blocking anti-TNF Ab Adalimumab, administered both subcutaneous and IA to the pregnant dams. We also tested their combined effect (see Fig. S9 for experimental scheme). Both reagents had previously been shown to be efficacious in macaques^22,23,35^.

Cytokine levels in the AW were only partially diminished with any of the treatments (Table S2). This modest effect was confirmed by the fact that cytokine mRNA expression in the lung was also minimally decreased in all the treatment groups (Table S3). LPS-induced histological changes (cell infiltration and lung interstitium thickening) remained present in animals given IL-1RA or anti-TNF, although lung interstitium thickening was partially decreased in animals given both inhibitors (Fig. 6A).

**Fig. 6.**
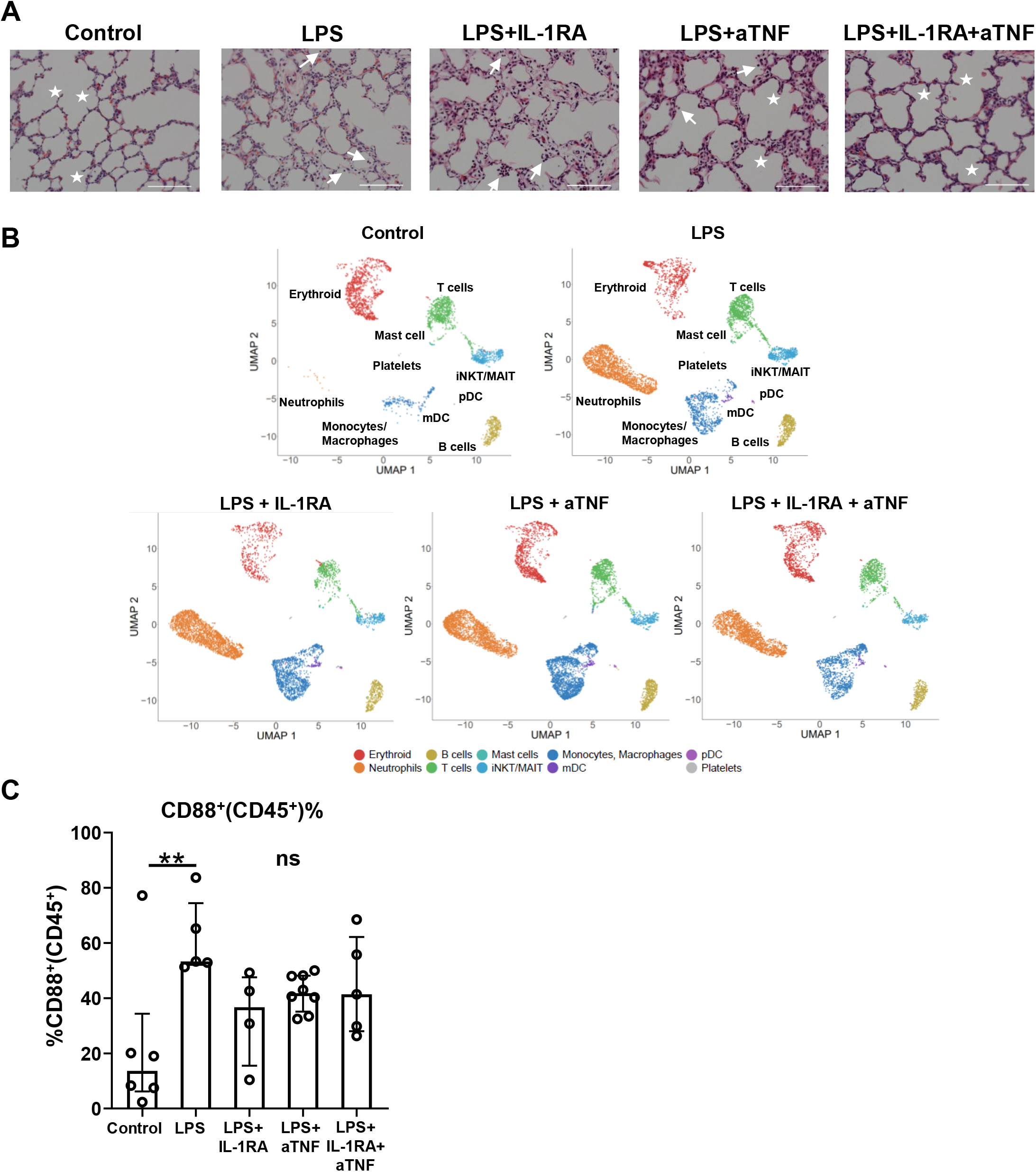
Hematopoietic cell changes in the fetal lung following blocking IL-1 and TNF signaling. **(A)** 20x H&E sections of fetal control, lps, lps+IL-1RA, lps+anti-TNF, and lps+IL-1RA+anti-TNF exposed animals; stars denote secondary septa, arrows represent cells in alveolar space and scale bar is 100μm **(B)** UMAPs of the hematopoietic cells from control, lps, lps+IL-1RA, lps+anti-TNF, and lps+ IL-1RA+anti TNF exposed animals. **(C)** CD88^+^ percentage within the CD45^+^ cells. Data presented as median an¢ interquartile range, Kruskai-Wallis test; **p≤0.01.

Infiltration of myeloid cells into the lung was not blunted by any of the treatments, as shown by both scRNAseq and flow cytometry analyses (Fig. 6B-C). Neutrophil accumulation in the AW was also not blunted (p>0.9, Kruskal-Wallis tests). Similar monocyte/macrophage populations were present in the lung of treated animals as in IA LPS animals (Fig. 7A). However, the blockades partially suppressed some of the biological processes activated by IA LPS (Fig. 7B and S10A-C). Notably, there was a downward trend of the expression of genes associated with the TLR, IL-1, and TNFα signaling pathways in the alveolar and interstitial macrophages, which appeared most marked in the animals that had received the combined treatment (Fig. 7B). Blockades also decreased the expression of genes associated with cell migration and apoptotic signaling in these cells (Fig. S10A-B). However, the blockades did not blunt these processes in the inflammatory monocytes (Fig. 7B and S10C).

**Fig. 7.**
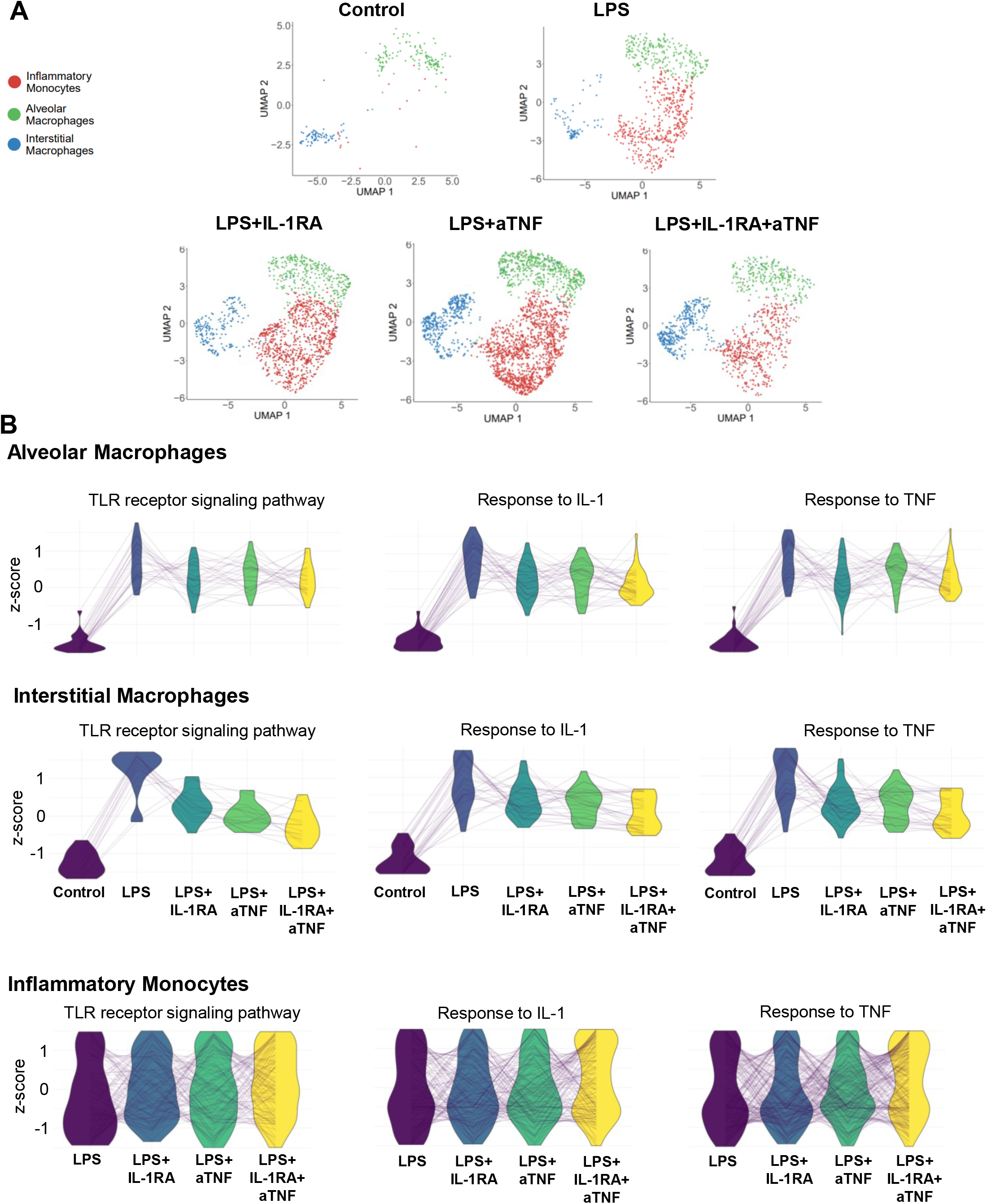
Fetal lung monocyte/macrophage population across treatment conditions. **(A)** UMAPs of fetal lung myeloid populations and **(B)** parallel coordinate plots of scaled expression of representative genes in select biological processes across treatment conditions in alveolar, interstitial macrophages and inflammatory monocytes.

Neutrophil migration into the lung was also not diminished by any of the blockades (Fig. 8A). As in the inflammatory monocytes, the blocking treatments did not markedly alter the neutrophilic transcriptome (Fig. 8B and S10D).

**Fig. 8.**
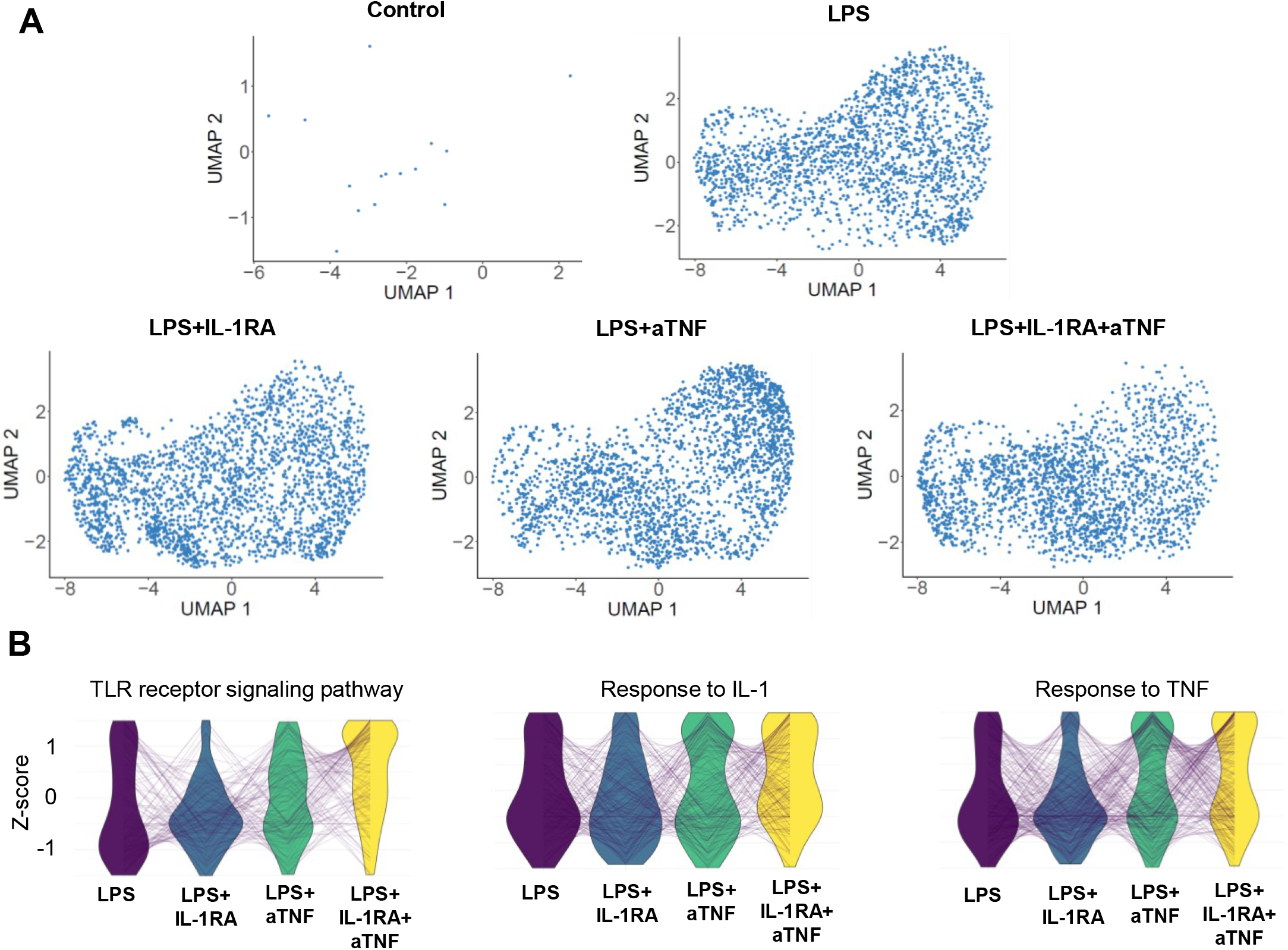
Fetal lung neutrophils in response to IA LPS and blockades. **(A)** UMAPs of the neutrophil population across treatment conditions. **(B)** Parallel coordinate plots of scaled expression of representative genes in select biological processes across treatment conditions.

Blockades lead to a decreased frequency of the mDC population, decreasing it below that present in the control animals (Fig. S11). mDC activation, as evidenced by high HLA-DR MFI, was diminished by the treatments (Fig. S11). In that case, the strongest effect was seen in animals that received IL-1RA, alone or in combination (Fig. S11).

## Discussion

Premature infants exposed to IUI are at increased risk of developing pulmonary and intestinal morbidities after birth ^9,36,37^. Herein, we applied several complementary techniques to obtain a comprehensive view of the fetal mucosal immune response to IUI in the highly relevant Rhesus macaque model.

In this model, shortly after IA LPS, robust inflammation developed in the fetal lungs, characterized by the accumulation of immune cells into the lung and increased levels of many inflammatory mediators both in the lung tissue and the AW, similar to what had previously been described in other IUI animal models ^10,11,28,29^. Myeloid cells were particularly involved, as the major cellular change occurring in the lungs of IA LPS-exposed fetuses was the massive accrual of neutrophils and inflammatory monocytes, which were absent in the control lungs. In addition, resident myeloid cells, both the alveolar and interstitial macrophages and the resident myeloid DCs, exhibited many signs of activation. Notably, scRNAseq analyses identified that response to pro-inflammatory cytokines, such as IL-1 and TNF, or to TLR, were activated in these subsets. Many of the activated biological processes were shared by the monocyte/macrophage and neutrophil populations, highlighting a general inflammatory transcriptional program. Overall, these data indicate that the fetal innate immune system can rapidly mount a strong response, a concept that remains controversial. Indeed, there have been many papers describing functional impairment in fetal/neonatal neutrophils and monocytes/macrophages as compared to their juvenile and adult counterparts (rev. in ^38–40^). Described defects notably include the decreased recruitment of neutrophils and inflammatory monocytes into the lung in response to *E. coli* instillation during the same time period, compared to juvenile mice ^41^. In contrast, our data in a NHP model argue that fetal neutrophils and inflammatory monocytes can quickly migrate into the pulmonary environment, as these two populations became very abundant in the lung 16hrs after IA LPS. Neutrophils appeared to localize in the lung in the form of aggregates, a pattern reminiscent of that described in murine models of acute lung injury ^42,43^. Similarly, other groups had described rapid recruitment of neutrophils in fetal sheep or rhesus macaque lungs in response to IA LPS or IL-1β ^27,28^. It is also to be noted that the concept of defective functionality of innate immune cells in human neonates mainly arises from studying *in vitro* stimulated cord blood cells. It is therefore possible that these assays do not fully reveal their potential. Another caveat to remember when interpreting data about preterm babies’ immune cell functions, is that many of these infants had just been given steroids, which affect various cell type functionality. We did not administer steroids in our studies, which may have allowed us to observe such robust immune responses.

Until recently, circulating and tissue neutrophils were considered as a homogeneous population with a defined array of functions. This concept has been challenged by many studies that revealed a high degree of variety in this population (rev. in ^44–49^). Two main mechanisms appear to drive this diversity, an intrinsic heterogeneity already initiated in bone marrow precursors on one hand, and on the other hand a highly dynamic transcriptional and epigenetic program triggered by exposure to the environment, notably in inflammatory conditions. Our pseudo-time analyses concur with the concept of functional heterogeneity, as they show several gene expression profiles within the neutrophils recruited to the lungs of IA LPS exposed animals. For example, genes related to granule content (*CAMP, LCN2*) or antimicrobial defense (*S100A8, S100A9*) were in general not simultaneously expressed along inflammatory mediators (*IL1A, IL1B, CCL2, CCL20*). Interestingly, *Bcl2A1*, a major pro-survival factor for neutrophils ^50^, including for LPS-activated neutrophils in the Rhesus chorio-decidua ^22^, was also heterogeneously expressed among the lung neutrophils in LPS-exposed animals, suggesting that differential survival may affect neutrophil heterogeneity. Future studies will be needed to further characterize at the protein level neutrophil heterogeneity and its impact on fetal responses to IUI.

We were also able to evaluate the role played by IL-1 or TNFα in fetal mucosal responses to IA LPS. Our data show that blocking either or both of these pathways had only a partial effect on lung inflammation, a finding consistent with the fact that fetal responses were likely driven by both direct activation by LPS and indirect cytokine-mediated pathways. The partial amelioration of disease by IL-1RA is similar to what has been previously described in other IUI models ^22,23,28,30^, while this was, to our knowledge, the first attempt at evaluating the effect of anti-TNF alone or in combination with IL-1RA. In general, the anti-TNF Ab appeared to have broader effectiveness than IL-1RA, which may be due to the fact that this treatment decreased levels of IL-1β expression in the lung tissue and AW, whereas IL-1RA did not affect TNFα levels (Tables S2-S3). Interestingly, the combination of both blockades did not radically ameliorate the effectiveness, which suggests that the two pathways are inter-connected, and that other mediators control the recruitment of myeloid cells into the lung. Notably, levels of CCL2, which regulates many aspects of monocyte biology, including recruitment, activation, and survival (rev. in ^51^), remained elevated in the lung tissue and AW of treated animals. Lung expression of CXCL12, which plays a role in myeloid cell recruitment during lung injury ^52,53^, might also be involved. Our data also show that high levels of IL-8 persisted in the lung and AW despite the blockades (Tables S2 and S3). Furthermore, many neutrophilic chemoattractant molecules, including chemotactic lipids, formyl peptides, and complement anaphylatoxins are also released during inflammation (rev. in ^54^). We did not quantify these mediators in fetal lungs or AW, but because they are released through both TLR and cytokine-mediated mechanisms, it is likely that they were still released despite the blockades. Interestingly, compared to the fetal lung, myeloid cell recruitment in the placenta was significantly reduced in the presence of either IL-1RA or anti-TNF ^22,35^, suggesting that the mechanisms controlling cellular migration might be tissue specific. Alternatively, and not exclusively, the fact that cells recruited in the placenta have a maternal origin, in contrast to fetal cells migrating to the lungs, may also have driven some of these differences.

Overall, our work in a model that phenocopies the main features of human IUI gives novel insights into the inflammation rapidly developing in the fetal lung following such exposure. One important conclusion from our new data is that the fetal immune system is highly responsive to its environment. In this model of acute sterile inflammation, we saw that fetal myeloid cells could be rapidly mobilized, migrating to the inflamed environment and displaying profiles of activation very similar to those described in adult cells. This confirms the new paradigm in the field of neonatal immunology, e.g., that fetuses have the capacity to mount a very robust innate immune response. Our data also emphasize the fact that the fetal lung is particularly responsive, as it is in close contact with the amniotic fluid, whereas the fetal gut appears less affected. This study thus provides more insights into why the most solid associations found between IUI exposure and post-natal morbidities in humans involved mainly the lung (bronchopulmonary dysplasia, wheeziness, and asthma). Our data also show that the multi-parametric mucosal inflammation that develops in IUI-exposed fetuses is not easily controlled by blocking only one, or two inflammatory mediators, emphasizing the need for additional mechanistic studies to better understand the dynamic and complex responses that IUI triggers in exposed fetuses.

## Materials and Methods

### Animals

The Institutional Animal Care and Use Committee at the University of California, Davis approved all animal procedures. Normally cycling, adult female Rhesus macaques (*Macaca mulatta*) were time mated. At approximately 130 days of gestation (about 80% of term gestation), the pregnant dams received either 1 ml saline solution or 1mg of LPS (Sigma-Aldrich, St. Louis MO) in 1 ml saline solution by ultrasound guided IA injection. IA administration of LPS or saline was performed in mothers of similar weights and ages with fetus with similar fetal genders and gestational ages (Table S1). Fetuses were surgically delivered 16, 48 or 120 hrs later by cesarean section. Delivered fetuses were euthanized with pentobarbital and tissues collected. There were no spontaneous deaths or preterm labor in the animals. Some pregnant dams were given, in addition to IA LPS, either the IL-1RA (Kineret^®^ Sobi, Stockholm, Sweden), or the anti-TNF Ab Adalimubab (HUMIRA^®^ AbbVie Chicago, IL), or both. These inhibitors were administered subcutaneously (100mg or 40mg for IL-1RA and anti-TNF respectively) 3hrs before IA LPS as well as IA (50mg or 40 mg for IL-1RA and anti-TNF respectively) 1hr before IA LPS (see scheme on Fig. S9).

### Histological evaluation of fetal lung and jejunum

Inflated lung (30cm water pressure) and jejunum tissues were fixed in 10% formalin immediately following removal. After a series of alcohol and xylene washes, tissues were blocked in paraffin. Paraffin sections were stained with hematoxylin & eosin (H&E). 20x images were taken with a Nikon 90i Upright Widefield Microscope (Nikon Instruments Inc., Melville, NY). 100μm scale bar added using Nikon Elements Software (Nikon Instruments Inc., Melville, NY).

### Immunofluorescence staining of neutrophils and confocal analysis

Upper left lung lobe sections (5μm thick) were blocked with 4 % goat serum in PBS, and then incubated 1hr at room temperature with the primary mouse-anti-human Ab diluted in 4% goat serum in PBS. Cross-reactivity of all Ab with Rhesus macaque cells was verified on adult tissues prior to their utilization to stain fetal cells. Ab were titrated for optimal detection of positive populations and MFI. Sections stained with unlabeled Ab were stained with a corresponding secondary goat-anti-mouse Ab conjugated with either Alexa Fluor 594, Alexa Fluor 488 or FITC (Life Technologies, dilution 1:400) for 1hr. As negative control, tissues were also stained with the secondary Ab only. Sections were washed and incubated with Vecta shield autofluorescence reduction kit and mounted with mounting media containing DAPI (Vector Laboratories). Images were acquired using an inverted microscope (Ti-E; Nikon, Japan) outfitted with a confocal scan-head (A1R; Nikon; Japan) at the CCHMC Confocal Imaging Core. Analyses involving deconvolution and background reduction were done with Nikon Elements Software.

Since neutrophils in the sections were mostly in clumps/aggregates, we evaluated the number of neutrophil aggregates in 10 fields per animal. The neutrophils were identified as S100A8-FITC (CF-145) and TNFAIP6-AF594 (polyclonal) (both ThermoFisher Waltham, MA) double positive cells. The average area covered by neutrophils clumps were also determined and normalized to the total DAPI area of the image per field. CD68-FITC (KP1) (Alligent Dako, Santa Clara, CA) and HLADR-AF594 (L243) (BioLegend, San Diego, CA) were also used to quantify neutrophils.

### Alveolar wash analyses

Alveolar wash was collected from the left lung. Supernatant was collected and frozen until cytokine analyses, using the NHP multiplex kit according to the manufacturer’s protocol (Millipore Austin, TX). Cell pellets was cytospun and slides were stained with Diff quick. 5 fields/slide (100 cells/field) were counted twice.

### Quantitative RT-PCR analyses of mucosal tissues

From frozen lung and jejunum tissue, total RNA was extracted after homogenizing in TRIzol (Invitrogen, Carlsbad, CA). A Nanodrop spectrophotometer (Thermo Fisher Scientific) was used to measure RNA concentration and quality. 1μg of RNA was used to generate cDNA using the Verso cDNA synthesis kit (Thermo Fisher Scientific), following the manufacturer’s protocol. Quantitative RT-PCR was done with a StepOnePlus real-time PCR system and rhesus specific TaqMan gene expression primers (Life Technologies, Carlsbad, CA). Eukaryotic 18S rRNA was the endogenous control used for normalization of the target RNAs. The mRNA fold change was calculated relative to the average value of the control group.

### Fetal lung cell isolation

The fetal lung was removed, and left lobe was processed into single cell suspensions as described previously ^55^. The large airways were removed and approximately ~500mg of airway free tissue was collected in a gentleMACS C-tube (Miltenyi Biotec, Auburn, CA) and 5mL digest buffer was added and placed on a gentleMACS octo dissociator for programmed runs. Following dissociation, the cell suspension was passed through a 100μm filter and washed with PBS. After washing, cells underwent red cell lysis (eBioscience, San Diego, CA) along with another pass through a 40um filter after neutralization.

### Single cell sequencing of fetal lung cells

scRNA-seq analyses were done in a subset of animals (Table S4). ~50,000 cells per lung were submitted for single cell sequencing at the Cincinnati Children’s Hospital Medical Center DNA Sequencing and Genotyping Core. Approximately ~16,000 cells were loaded into one channel of the Chromium system using the 3 prime v3 single cell reagent kit (10X Genomics, Pleasanton, CA). Following capture and lysis, cDNA was synthesized and amplified as per the manufacturer’s protocol (10X Genomics). The amplified cDNA was used to construct Illumina sequencing libraries that were each sequenced using an Illumina HiSeq 4000 machine. Raw sequencing data was processed aligned to the Rhesus macaque reference Mmul_8.0.1 (Ensembl version 91) with Cell Ranger 3.0.2 (10X Genomics) generating expression count matrix files.

After doublet removal with the software DoubletFinder (version 2.0), an integrated analysis was performed using R (version 3.6.3) and the Seurat package (version 3.1.0, https://satijalab.org/seurat/), to identify common cell types across different experimental conditions. Specifically, cells with 25% or more mitochondrial transcripts were removed, as well as cells expressing fewer than 150 or more than 5000 features. After log-normalization, 5000 highly variable features were identified per sample using the vst method. Integration was performed using the Seurat FindIntegrationAnchors and IntegrateData functions with default parameter values. The integrated data was scaled, including regressing out mitochondrial percentage and cell cycle variables, followed by application of PCA reduction. A UMAP projection was estimated based on the PCA reduction and the correlation metric. Clustering of cells was done using the Seurat functions FindNeighbors (based on first 30 PCs) and FindClusters (resolution=4). This high-resolution was initially used to define highly discrete immune populations, with the final populations subsequently aggregated into broader, higher confidence cell-types based common marker genes expressed and corresponding UMAP embeddings. To annotate these cell-populations, were performed a gene-set enrichment analysis on the aggregated cell populations used established human and mouse cell-population marker genes annotated by LungMAP, from the AltAnalyze and ToppCell databases. Immune cell population labels were further evidenced using label transfer with the software cellHarmony against a recently described reference human lung datasets^56,57^. Clusters were orthogonally validated using the software ICGS2 in AltAnalyze, which does not require a resolution setting, to confirm their presence. Identified immune cell clusters were subset into a new Seurat object and re-clustered. For further analyses, Y-chromosomal genes were removed from the dataset because both IA LPS-exposed fetuses were male whereas animals of both sexes were present in the other experimental groups. Additionally, we checked for the expression of genes *XIST* and *TSIX*, which did not appear in our dataset and seem to not be drivers of differences observed. All identified cell populations were required to have unique reported marker genes to ensure they were not artifacts of “over-clustering”. Differential gene expression analysis was performed as implemented in Seurat. Plots were generated using the ggplot2 package (version 3.3.2). Differentially expressed genes (DEGs) between control and LPS in the monocyte/macrophage cluster with a fold change (FC) of ≥ 1.2 and adjusted p-value of ≤ 0.05 were used for functional enrichment analysis of biological processes and pathways using the ToppFun web portal^58^. Terms with higher expression in LPS are represented as positive log p-values. Scaled expression of genes for upregulated biological processes terms in the monocyte/macrophage cluster were used to plot parallel coordinate plots using the ggplot2 package.

Differential gene expression analysis was not feasible for the neutrophil or inflammatory monocyte clusters since those cells were not present in control samples (<20 cells). We thus performed single sample gene set enrichment analysis (ssGSEA) in R using the ssgsea2.0 package (github.com/broadinstitute/ssGSEA2.0) in order to identify enriched gene ontology (GO) gene sets (MSigDB v7.0). Expression values of overlapping genes of the most significant and relevant gene sets were plotted as parallel coordinate plots using ggplot2.

In order to assess possibly different states of gene expression within neutrophils after LPS exposure, single-cell trajectory analysis was performed with Monocle3 (version 0.2.3.0). Spatial autocorrelation analysis as implemented in Monocle3 was used to determine genes that most strongly vary along the pseudo-time trajectory. The Monocle algorithm is capable of ordering cells based on their transcriptomic profile in an unsupervised manner and thus arrange cells along a directional path. The order of the cells along this path represents different transcriptomic states within a biological process even if cells are obtained at only one time point during an experiment. The scRNA-seq data have been deposited in the Gene Expression Omnibus (GSE169390, reviewer token:qxatqgkyhxyrhkl).

### Flow Cytometry

A cocktail of conjugated Ab was used to phenotype lung cell suspensions for multi-parameter flow-cytometry. The following Ab were used: HLA-DR (L243), CD11c (3.9) (BioLegend, San Diego, CA); CD3 (GHI/61), CD3 (SP34-2), CD19 (SJ25C1), CD20 (2H7), CD123 (7G3), CD45 (DO58-1283) (BD Biosciences, San Jose, CA); CD8α (RPA-T8), FoxP3 (PCH101), LIVE/DEAD Fixable Aqua dead cell stain, (eBioscience, San Diego, CA), CD88 (P12/1) (Bio-rad, Hercules, CA). Lung cell suspensions were treated with human IgG to block Fc-receptors and stained for surface markers. Cells were then washed and treated with fixation/permeabilization buffer (ThermoFisher). Following permeabilization, cells were stained for intracellular markers, washed, and resuspended in PBS. Cells were collected on a FACS Fortessa (BD Bioscience San Jose, CA). BD FACSDiva Software v8.0.1 (BD Bioscience San Jose, CA) was used for analysis.

### Statistical analyses

GraphPad Prism 8 (GraphPad Software, La Jolla, California, USA) was used to graph and analyze data for statistical significance. Data were first checked for normality and values expressed as either mean ±SEM or median and interquartile range. Statistical differences between 2 groups were analyzed using Mann-Whitney U-tests or Student’s t-test. For comparison of more than two groups, Kruskal-Wallis or One-way ANOVA were used. Results were considered significantly different for p values ≤0.05. However, due to the limited number of samples in some groups, we also report trends (p values between 0.05 and 0.1).

## Author contributions

C.A.C, A.H.J., H.D., S.G.K., L.A.M., I.P.J. and W.J.Z. conceived the study. C.M.J., S.M., P.P., T.I., P.S., M.C., M.D. and J.G. all performed experiments and data analysis. N.S. and K.C. did the initial pre-processing of scRNAseq 10X files. All authors critically reviewed the manuscript and approved the final version.

## Acknowledgments

The authors would like to thank the staff at the California National Primate Research Center for their outstanding technical support, and especially Paul-Michael Sosa, Jennifer Kendrick, and Sarah Davis and for invaluable help in animal management and care. We also thank Cincinnati Children’s Hospital Medical Center’s Research Flow Cytometry, Confocal Imaging, and DNA Sequencing and Genotyping Cores.

