## Supplemental Figures, Tables 1-4 for "A potent myeloid response is rapidly activated in the lungs of premature Rhesus macaques exposed to intra-uterine inflammation"

Fig. S1

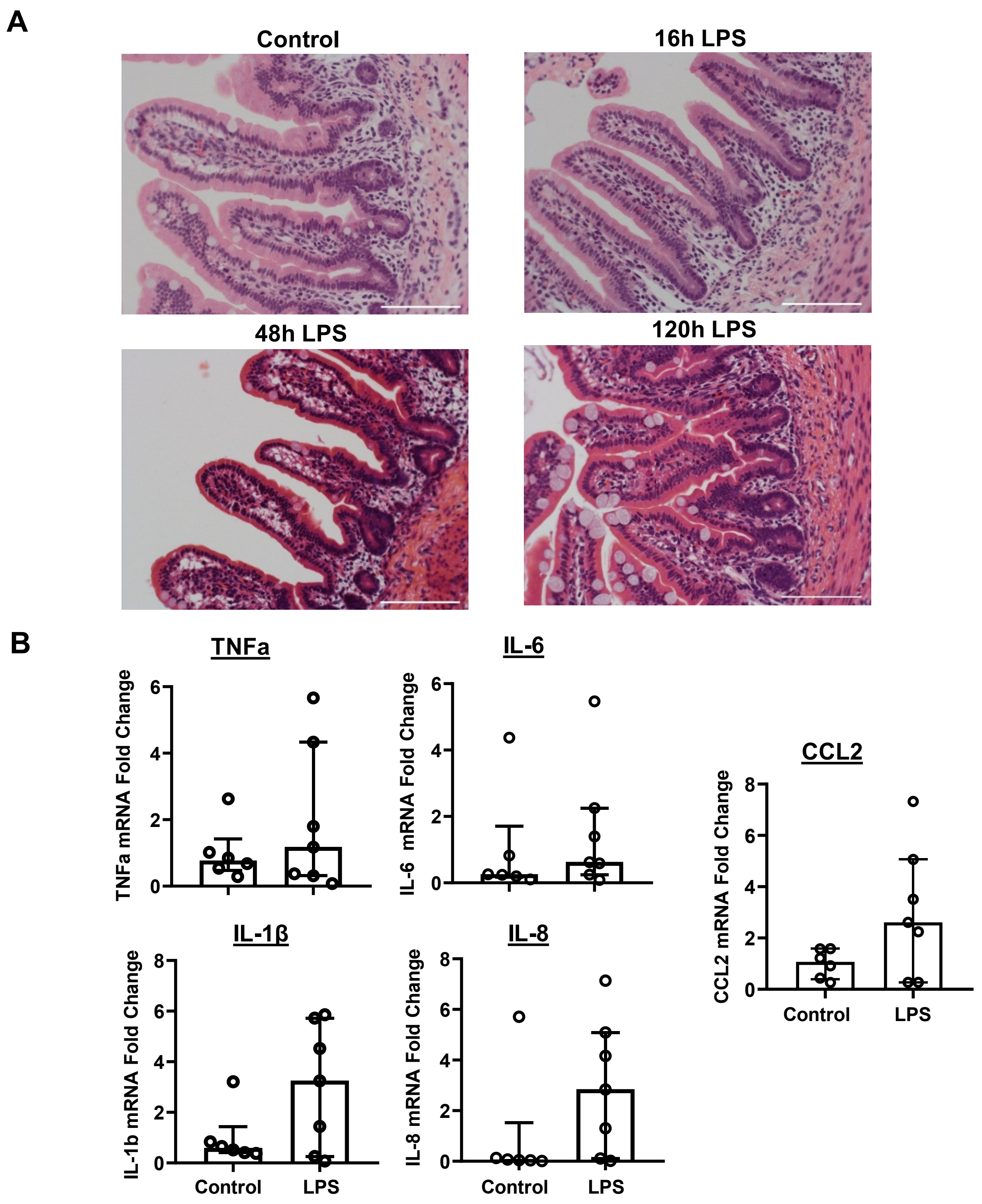


**Fig. S1. Jejunum histology and cytokine mRNA expression following IA LPS. (A)** Representative 20x H&E sections of fetal jejunum in control (upper left, n=4) and IA LPS, either 16 (upper right, n=5), 48 (bottom left, n=3), or 120 hours (bottom right, n=3) later; scale bar is 100μm. **(B)** TNFα, IL-6, IL-1β, IL-8, and CCL2 mRNA expression in the jejunum 16 hours post intra-amniotic LPS injection. Data presented as mean with SEM, student's unpaired t-test (IL-8) and median and interquartile range, Mann-Whitney U test (IL-6, TNFα, IL-1β, CCL2).

Fig. S2

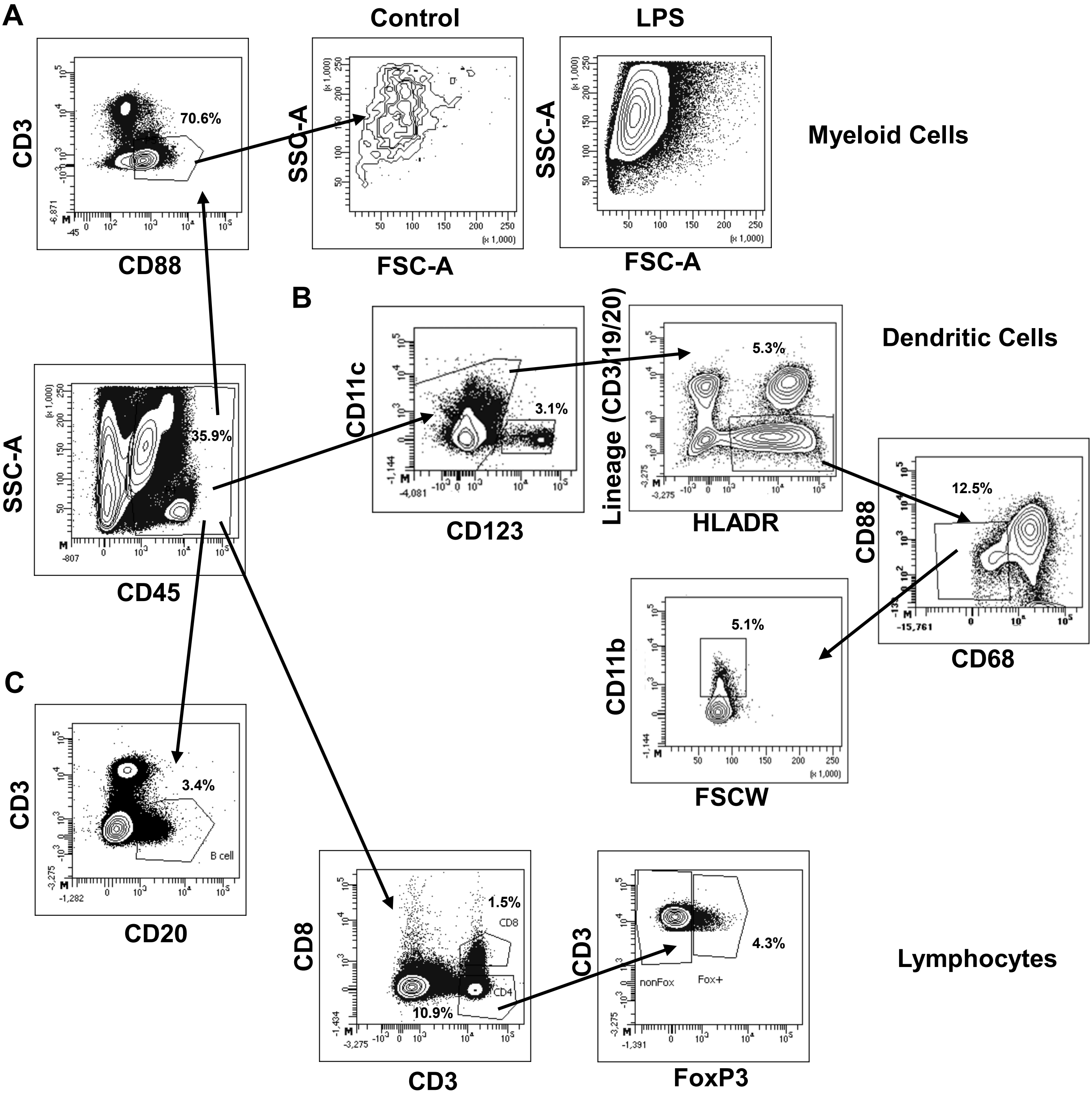


**Fig. S2. Flow cytometry gating strategy to identify immune cell populations in the fetal lung.** Representative flow cytometry gating of unstimulated lung single cells suspensions from a LPS-exposed fetus stained with a series of antibodies to identify **(A)** myeloid cells ($CD45^+CD88^+$), **(B)** dendritic cells populations identified were pDC ($CD123^+CD45^+$), mDC ($CD11c^+Lineage^-HLA\text{-}DR^+CD88^-CD68^-CD11b^+$). **(C)** Lymphocyte populations $CD4^+$ ($CD3^+CD8^-$), $CD8^+$ ($CD3^+CD8^+$), regulatory T cells ($CD3^+CD8^-FoxP3^+$) and B cells ($CD20^+$).

**Fig. S3**

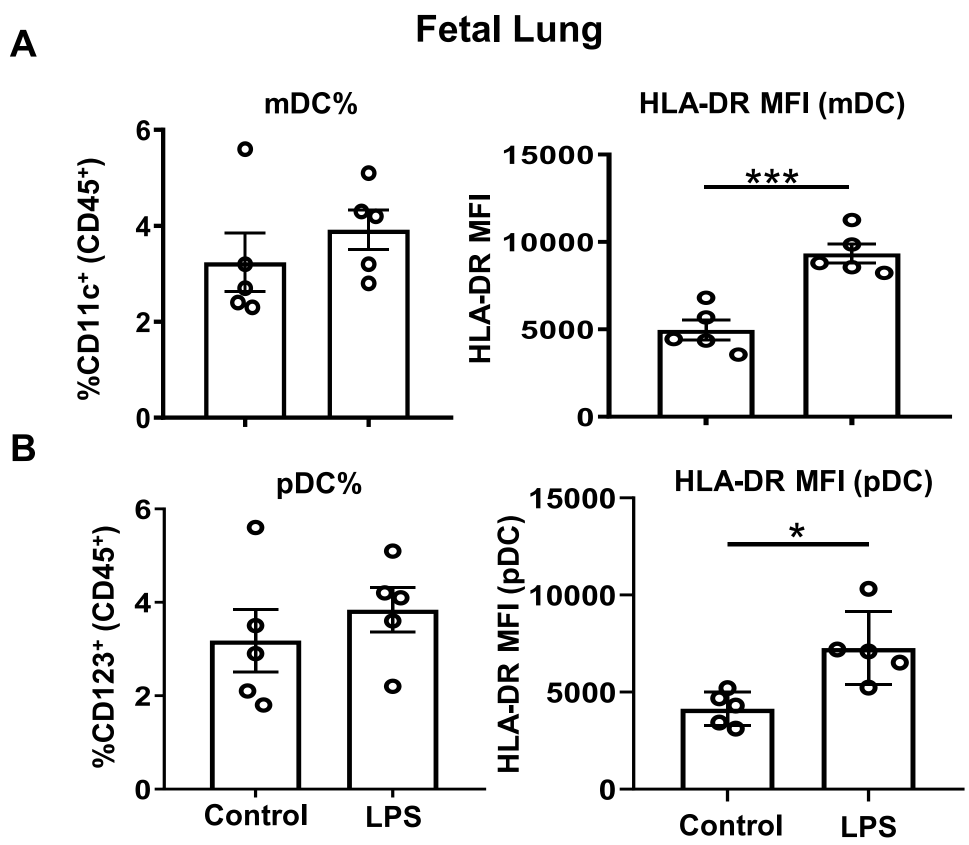


**Fig. S3. Dendritic cell response in fetal lung following IA LPS.** Percentage and HLA-DR MFI of **(A)** mDC (left and right) and **(B)** pDC (left and right) cells. Data presented as mean with SEM, student's unpaired t-test; $^{*}p \leq 0.05$, $^{***}p \leq 0.001$.

Fig. S4

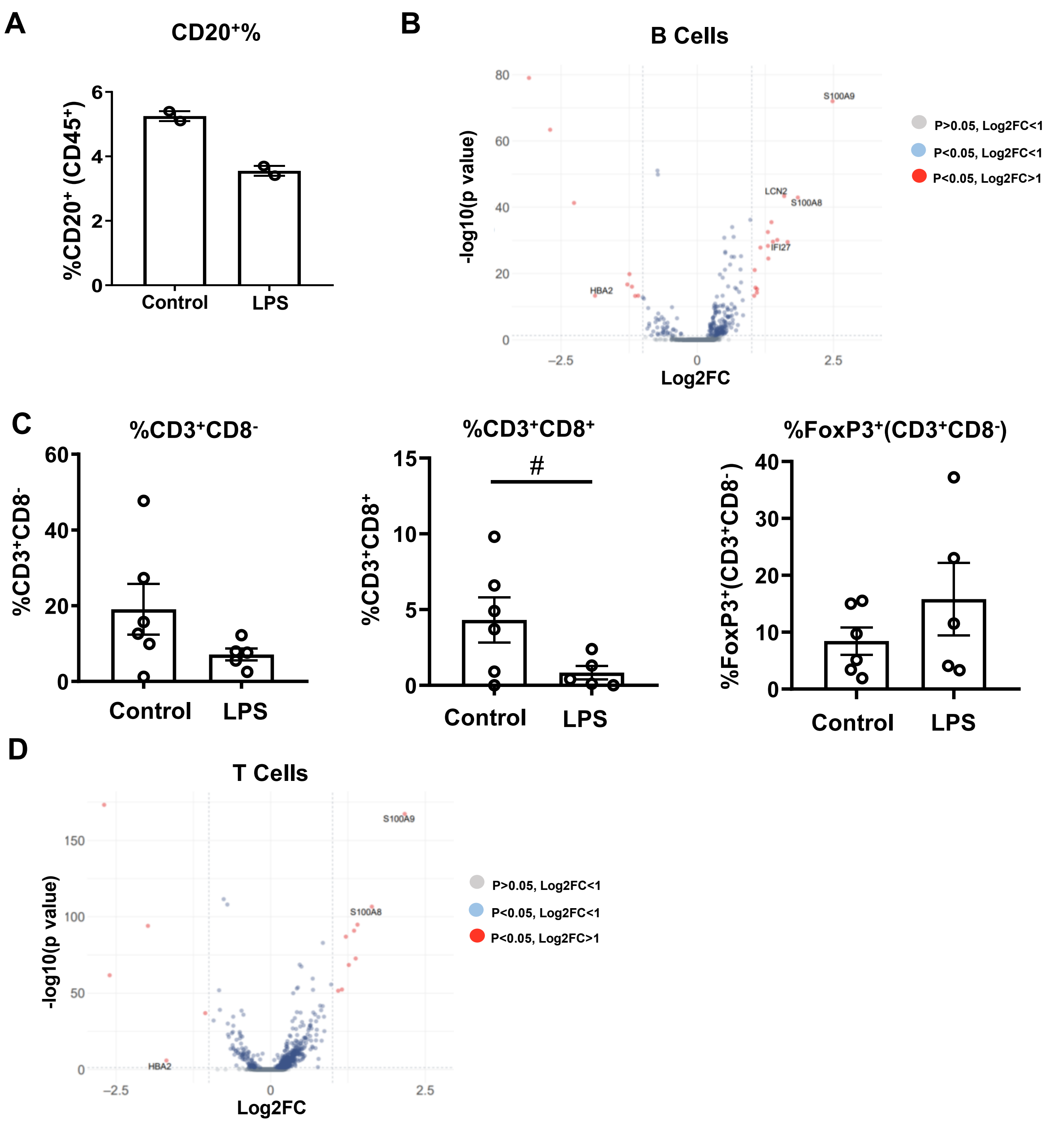


**Fig. S4. Lymphoid cell changes in fetal lung following IA LPS. (A)** Percentage and **(B)** volcano plot of differentially expressed genes in the B cells ($CD20^+$) population in control and LPS-exposed animals at 16 hours post IA injection of LPS. **(C)** Percentage of $CD4^+$($CD3^+CD8^-$), $CD8^+$, and $FoxP3^+$ T cells and **(D)** volcano plot of total T cell population in control and LPS-exposed animals. Data presented as mean with SEM, student's unpaired t-test (panel C) or median and interquartile range, Mann-Whitney U test (panel A); #p0.05≥x≤0.10.

### Fig. S5

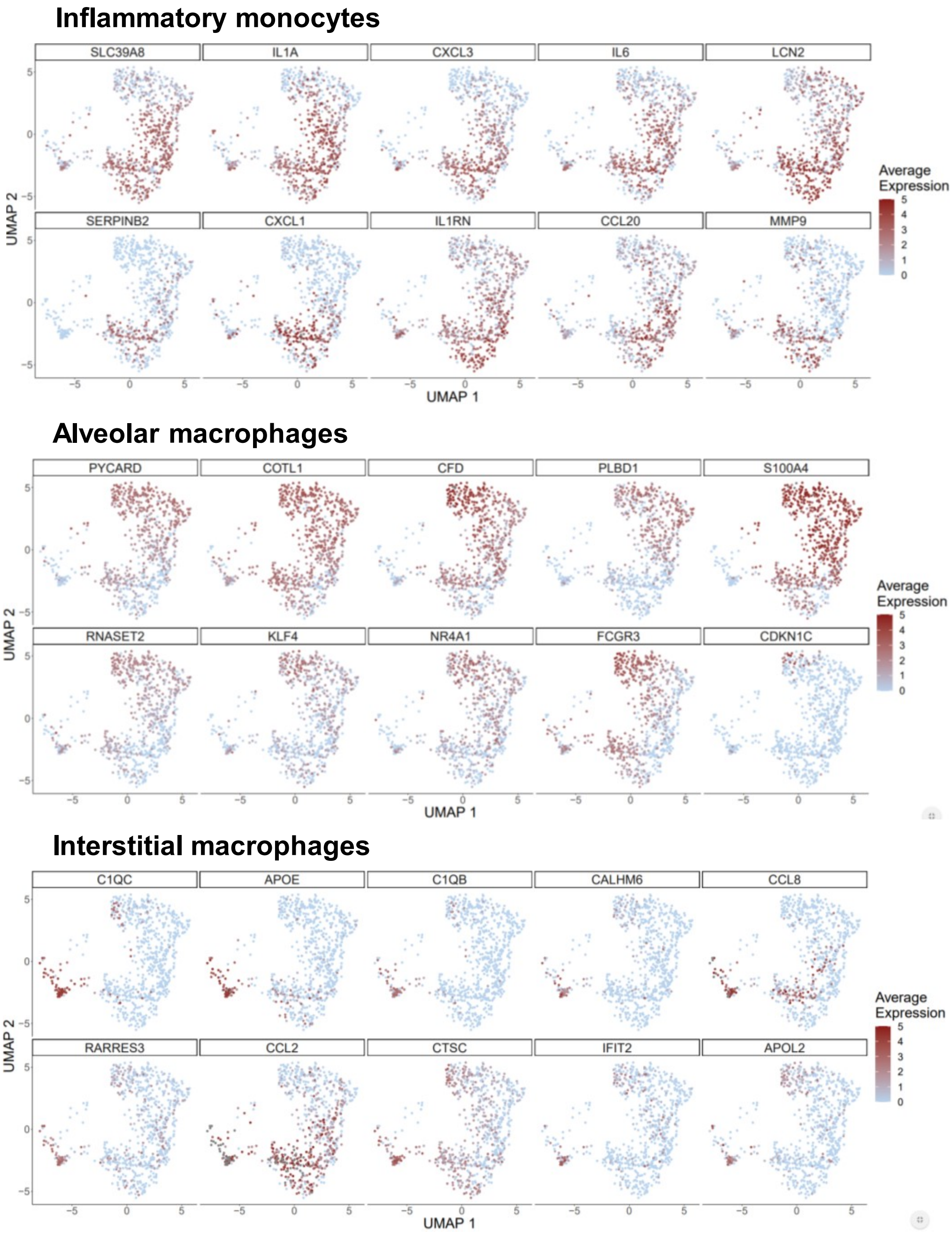


**Fig. S5. Differentially expressed genes in fetal lung monocyte/macrophage populations.** Feature plots of the top 10 conserved genes that were not changed between treatment groups in the inflammatory monocyte, alveolar macrophage, and interstitial macrophage populations in the fetal lung of IA LPS exposed fetuses.

**Fig. S6**

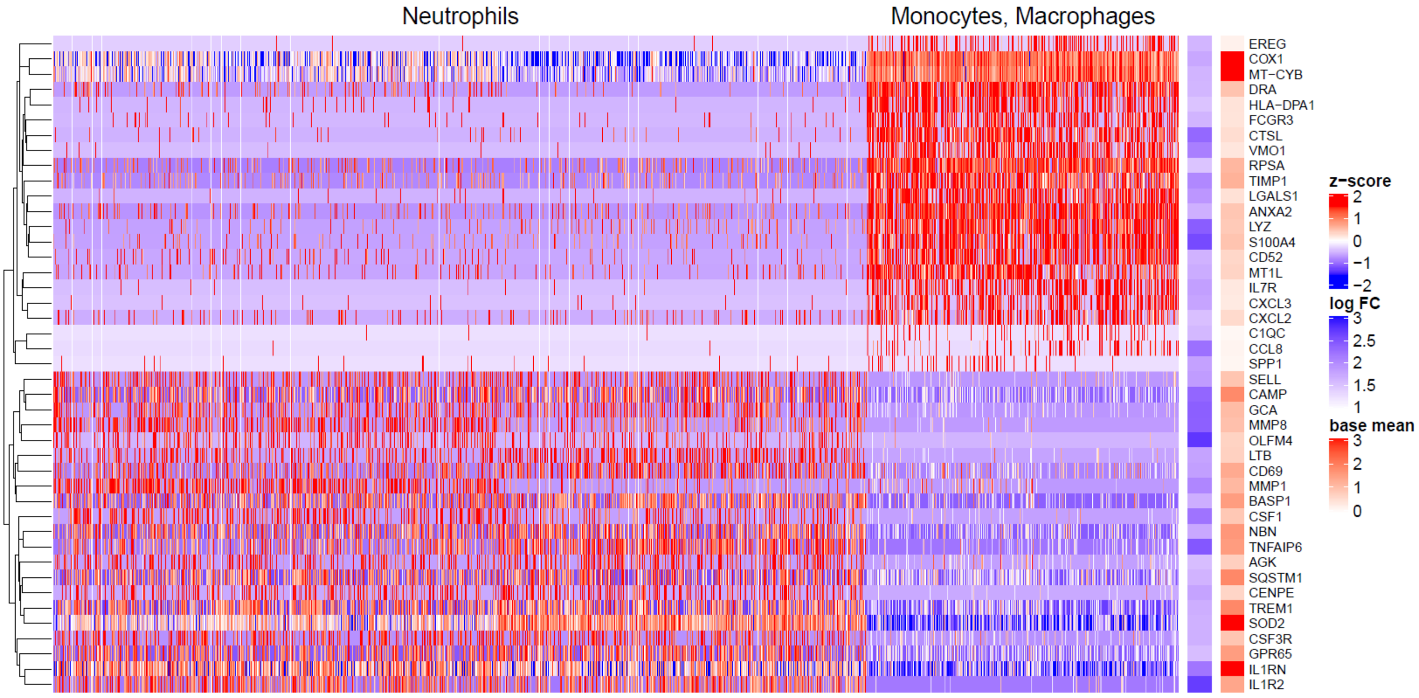


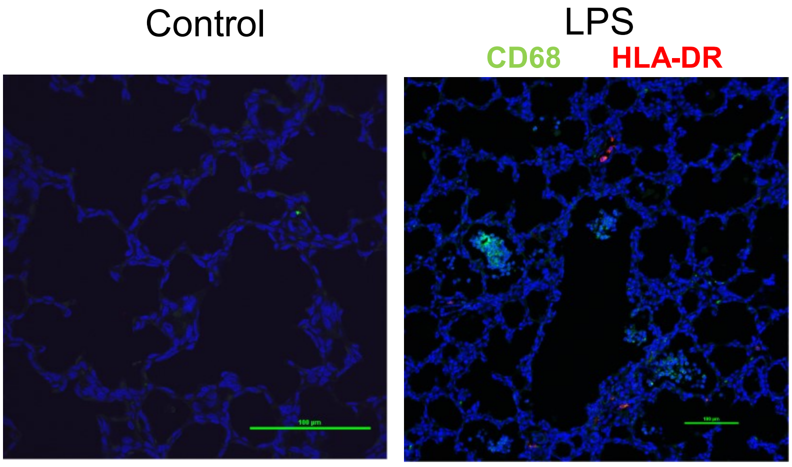


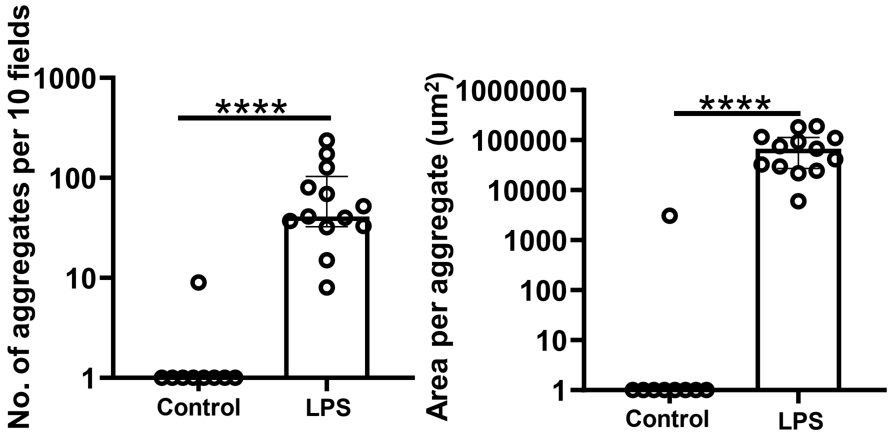


**Fig. S6. Myeloid populations in the fetal lung of IA LPS exposed animals. (A)** Heat map of top 43 genes (fold change ≥1.5) in neutrophils and monocytes/macrophages in the fetal lung of IA LPS exposed animals. **(B)** Immunofluorescence of IA control and LPS fetal lung stained for CD68 and HLA-DR; scale bar is 100µm. **(C)** Neutrophil aggregate count (left) and area (right) in the fetal lungs of control and IA LPS fetuses. Data presented as median and interquartile range, Mann-Whitney U test; ****p≤0.0001.

### Fig. S7

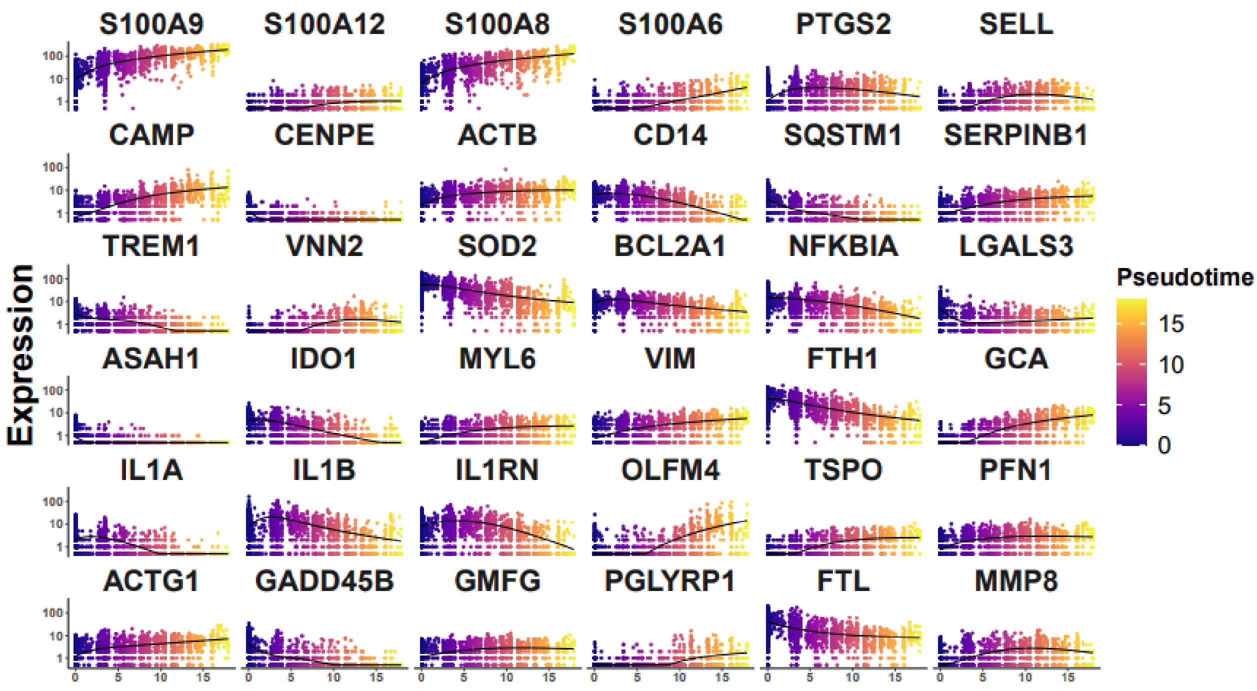


**Fig. S7. Pseudo-time analysis fetal lung neutrophils following IA LPS exposure.** Top 36 genes in fetal lung neutrophils that most drive the pseudo-time trajectory.

### Fig. S8

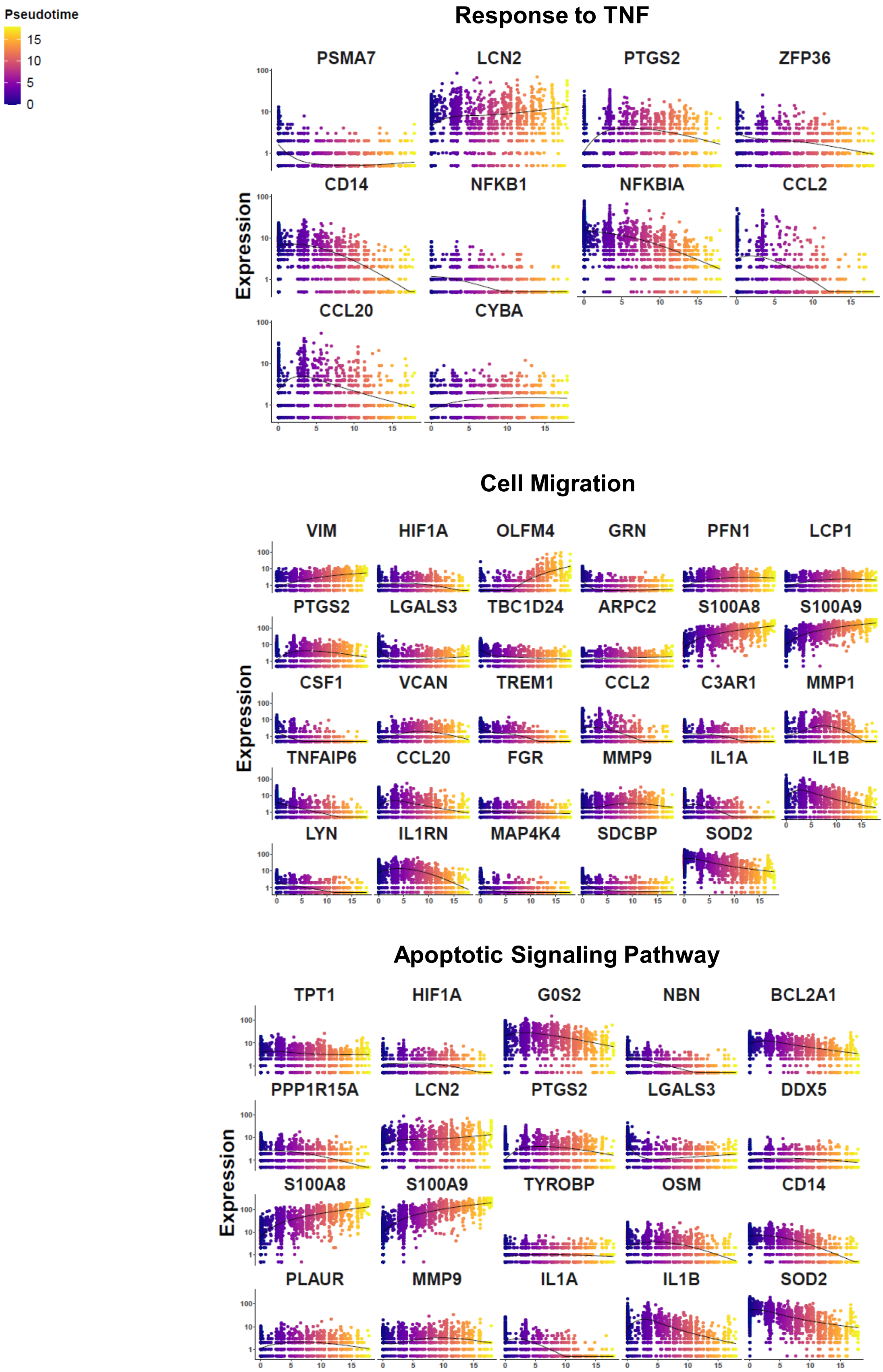


**Fig. S8. GO analysis biological pathways in IA LPS exposed fetal lung neutrophils.** Gene expression in fetal lung neutrophils across pseudo-time.

Fig. S9

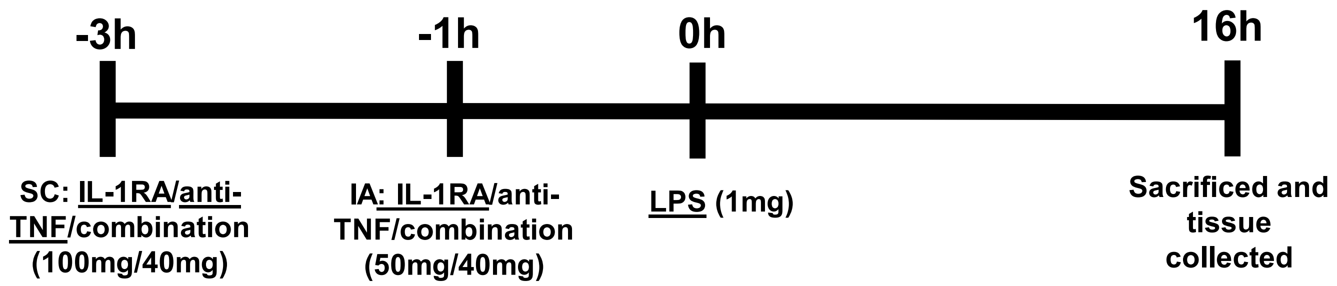


**Fig. S9. Fetal lung following IA LPS with blockades.** In blocking studies, mothers received either IL-1RA (Kineret), anti-TNF (adalimumab), or both 3 (subcutaneous) and 1 hour (IA) before IA LPS.

### Fig. S10

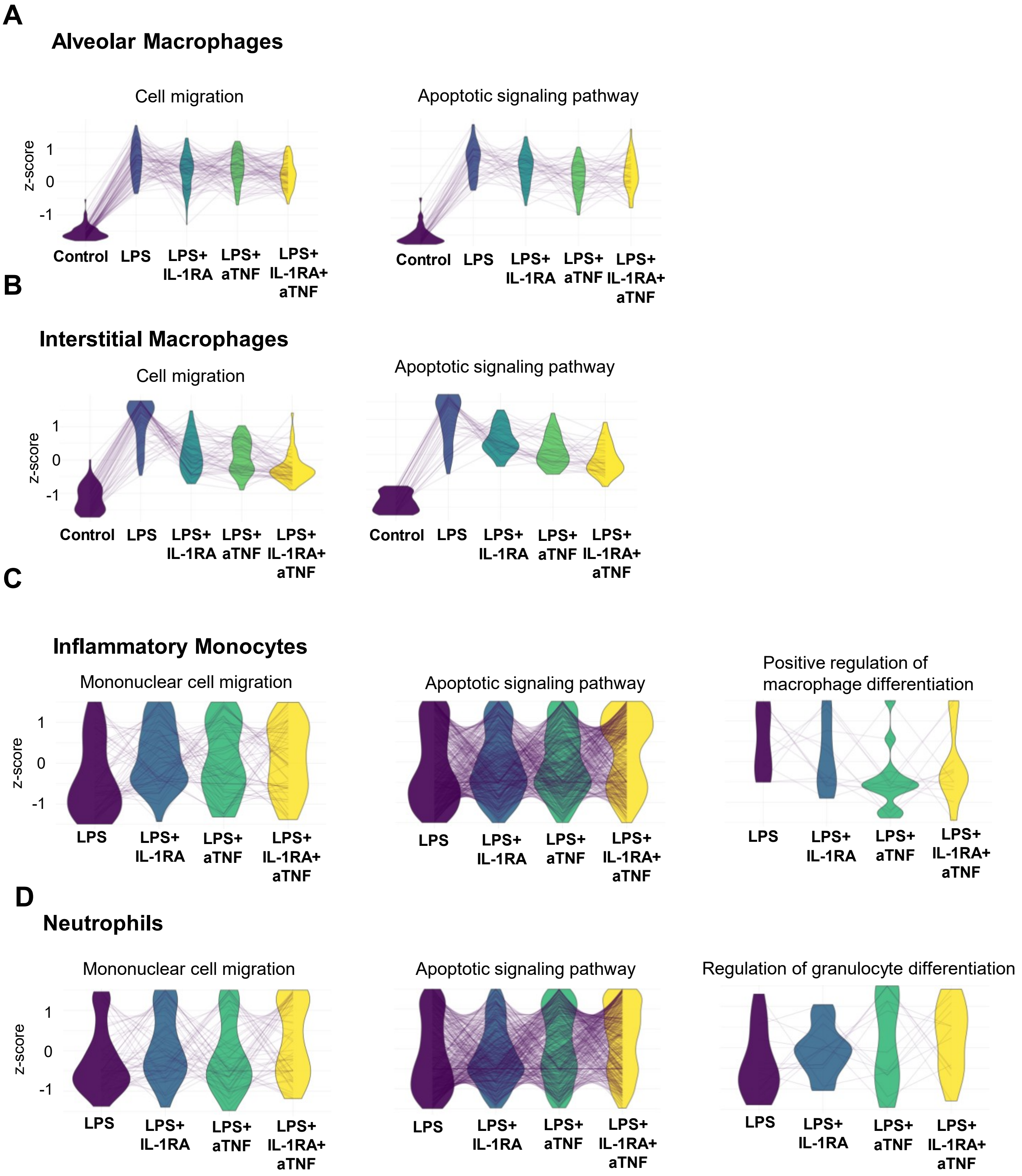


**Fig. S10. Transcriptional profile of myeloid populations in the fetal lung across treatment conditions.** Parallel coordinate plots of scaled expression of representative genes in select biological processes across treatment conditions in the **(A-C)** monocyte/macrophage population and **(D)** neutrophils.

Fig. S11

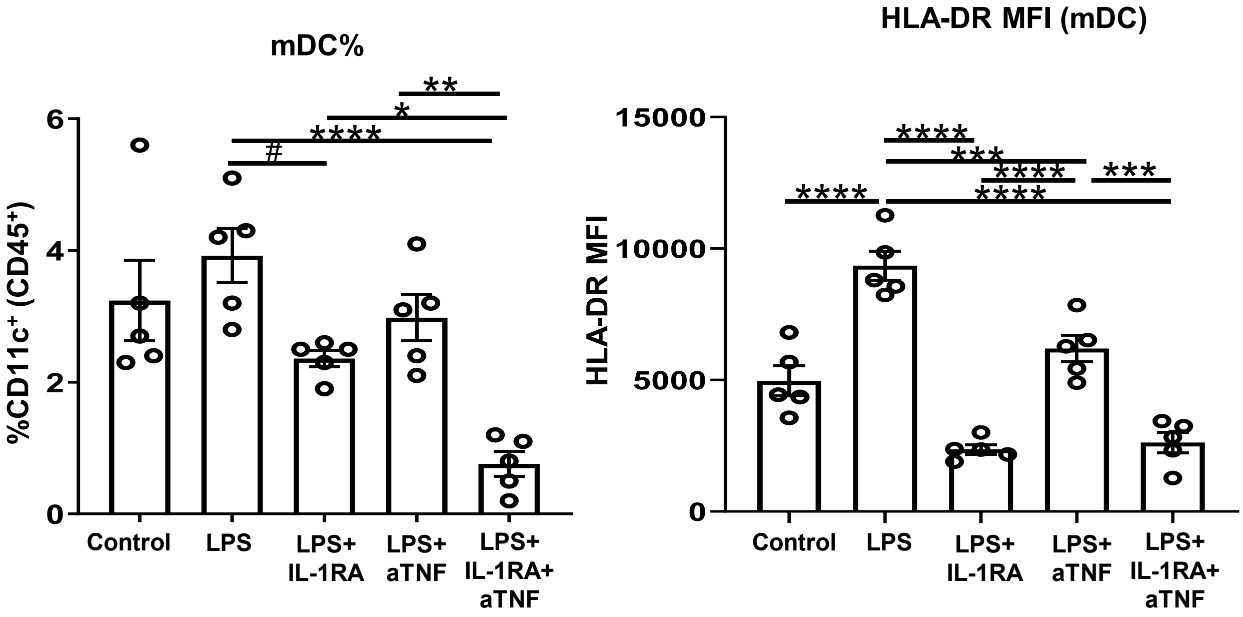


**Fig. S11. mDC changes in fetal lung following blocking IL-1 and TNF signaling.** Percentage and HLA-DR MFI of mDCs (left and right). Data presented as mean with SEM, one-way ANOA (mDC% and HLA-DR MFI) $^{+}$%; *p≤0.05, **p≤0.01, ***p≤0.001, ****p≤0.0001, and #p0.05≥x≤0.10.

**Table S1. Demographic data of fetal animal used in the study**

| | *Controls (n=24) | LPS (16hr) (n=21) | LPS+IL-1RA (n=10) | LPS+aTNF (n=14) | LPS+IL-1RA+ aTNF (n=5) |
|---|---|---|---|---|---|
| Fetal gestational age (days) | 132±0.5 | 132±0.6 | 132±0.9 | 131±1.0 | 133±0.8 |
| Fetal birth weight (g) | 330±5.9 | 330±9.5 | 338±13.3 | 355±19.4 | 359±17.3 |
| Fetal sex (%female) | 33 | 67 | 60 | 43 | 20 |

mean±SEM; *One control animal is missing gestational age

**Table S2. Cytokine production in alveolar wash**.

| ng/mL | Controls (n=20-24) | LPS (16hr) (n=19-21) | LPS+IL-1RA (n=8-10) | LPS+aTNF (n=13-14) | LPS+IL-1RA+ aTNF (n=4-5) |
|---|---|---|---|---|---|
| TNFα*+ | 0.0038 (0.0001-0.021) | 0.448 (0.0316-3.713) | 0.825 (0.0311-4.595) | 0.052 (0.0036-0.097) | 0.039 (0.030-0.059) |
| IL-6* | 0.295 (0.0042-2.677) | 3.148 (0.330-17.40) | 1.293 (0.114-3.253) | 1.422 (0.153-7.365) | 1.146 (0.293-2.802) |
| IL1β* | 0.00062 (0.0001-0.001) | 0.464 (0.0042-4.546) | 0.332 (0.003-0.696) | 0.151 (0.0030-1.312) | 0.055 (0.016-0.147) |
| GM-CSF* | 0.0017 (0.0004-0.009) | 0.539 (0.059-2.540) | 0.189 (0.118-0.275) | 0.369 (0.0098-2.257) | 0.107 (0.021-0.233) |
| IL-8* | 0.0976 (0.0072-0.708) | 18.58 (1.963-110.4) | 14.50 (2.309-15.72) | 5.642 (0.4417-12.53) | 4.281 (1.604-9.286) |
| CCL2* | 0.173 (0.0402-0.507) | 20.48 (0.0073-32.83) | 5.069 (1.248-10.31) | 8.832 (0.854-48.90) | 4.665 (1.357-10.85) |
| IL-10* | 0.0026 (0.0009-0.009) | 0.068 (0.0036-0.190) | 0.018 (0.0052-0.066) | 0.039 (0.001-0.205) | 0.062 (0.0094-0.109) |

Data presented as mean and range, comparisons made using Kruskai-Wallis test; *control v. lps $p \leq 0.0001$, + lps v. lps+aTNF $p \leq 0.05$

**Table S3. Cytokine mRNA expression in the fetal lung..**

| mRNA relative expression | Controls (n=18) | LPS (16hr) (n=15) | LPS+IL-1RA (n=6-9) | LPS+aTNF (n=6-9) | LPS+IL-1RA+ aTNF (n=5) |
|---|---|---|---|---|---|
| TNFα* | 1 (0.30-4.5) | 179 (24.1-421.2) | 186 (14.4-369.1) | 80 (9.9-153.6) | 165 (16.4-581.5) |
| IL-6* | 1 (0.53-2.2) | 1381 (128.2-2977.8) | 621 (60.2-963.5) | 458(18.6-1311.9) | 157 (26.2) |
| IL1β* | 1 (0.57-1.8) | 1021 (191.1-3039.4) | 984 (45.9-1403.3) | 423(48.2-790.9) | 420 (79.0-1256.1) |
| IL-8* | 1 (0.33-2.7) | 2993 (329.8-8800.6) | 3377 (159.3-8663.4) | 827 (58.1-2120.9) | 753 (85.4-2815.9) |
| CCL2* | 1 (0.02-2.5) | 138 (11.6-606.0) | 90 (9.7-191.5) | 43 (3.1-142-9) | 91 (15.9-284.4) |

Data presented mean and range, comparisons made using Kruskai-Wallis test; *control v. lps $p \leq 0.0001$

**Table S4. Demographic data of fetal animals used for scRNAseq**

| | Controls (n=2) | LPS (16hr) (n=2) | LPS+IL-1RA (n=3) | LPS+aTNF (n=3) | LPS+IL-1RA+ aTNF (n=3) |
|---|---|---|---|---|---|
| *Fetal gestational age (days) | 129.5±0.5 | 130.5±0.5 | 131.7±0.7 | 134.00±0 | 133.0±1.2 |
| *Fetal birth weight (g) | 348±48.0 | 298.4±38.7 | 301.7±26.1 | 416.4±32.7 | 343.1±12.5 |
| Fetal sex (%female) | 50 | 100 | 66.6 | 33.3 | 33.3 |

* Values are in mean±SEM
